# Arabidopsis glutathione reductase 2 is indispensable in plastids, while mitochondrial glutathione is safeguarded by additional reduction and transport systems

**DOI:** 10.1101/610477

**Authors:** Laurent Marty, Daniela Bausewein, Christopher Müller, Sajid Ali Khan Bangash, Anna Moseler, Markus Schwarzländer, Stefanie J. Müller-Schüssele, Bernd Zechmann, Christophe Riondet, Janneke Balk, Markus Wirtz, Rüdiger Hell, Jean-Philippe Reichheld, Andreas J. Meyer

## Abstract

- A highly negative glutathione redox potential (*E*_GSH_) is maintained in the cytosol, plastids and mitochondria of plant cells to support fundamental processes, including antioxidant defence, redox regulation and iron-sulfur cluster biogenesis. Out of two glutathione reductase (GR) proteins in Arabidopsis, GR2 is predicted to be dual-targeted to plastids and mitochondria, but its differential roles in these organelles remain unclear.
- We dissected the role of GR2 in organelle glutathione redox homeostasis and plant development using a combination of genetic complementation and stacked mutants, biochemical activity studies, immunogold labelling and *in vivo* biosensing.
- Our data demonstrate that GR2 is dual-targeted to plastids and mitochondria, but embryo lethality of *gr2* null mutants is caused specifically in plastids. Whereas lack of mitochondrial GR2 leads to a partially oxidised glutathione pool in the matrix, the ABC transporter ATM3 and the mitochondrial thioredoxin system provide functional backup and maintain plant viability.
- We identify GR2 as essential in the plastid stroma, where it counters GSSG accumulation and developmental arrest. By contrast a functional triad of GR2, ATM3 and the thioredoxin system in the mitochondria provides resilience to excessive glutathione oxidation.

## Introduction

The use of oxygen by aerobic organisms allows them to obtain more energy from carbohydrates, by accessing a larger reduction potential difference. Other metabolic reactions in the cell, however, also lead to reduction of oxygen and give rise to reactive oxygen species (ROS), such as superoxide (O_2_^−^), hydrogen peroxide (H_2_O_2_) and hydroxyl radicals. Low levels of ROS have been implicated in signalling between subcellular compartments as well as long-distance signalling regulating plant development and metabolism (Foyer & Noctor, 2009; Gilroy *et al*., 2016). However, at higher concentrations ROS may oxidize lipids, proteins and DNA and render these molecules non-functional. Therefore, ROS constitute a major threat to the cell and need to be tightly controlled. Cellular compartments that can be particularly affected by oxidative stress include the plastids, mitochondria and peroxisomes (Mittler *et al*., 2004). In chloroplasts, the Mehler reaction and antenna pigments are major sources of O_2_^−^ formation (Asada, 1999). In mitochondria, over-reduction of the electron transport chain results in O_2_^−^ production at complexes I and III (Moller, 2001). In both compartments, O_2_^−^ quickly dismutates to H_2_O_2_ and O_2_ in a reaction catalysed by superoxide dismutase (SOD).

In plants, H_2_O_2_ produced by SODs is further reduced to water by peroxidases such as ascorbate peroxidases (APXs), glutathione peroxidase-like enzymes (GPXLs) and peroxiredoxins (PRXs). While GPXLs and PRXs are dependent on thioredoxins (TRXs) as primary electron donor (Finkemeier *et al*., 2005; Navrot *et al*., 2006; Attacha *et al*., 2017), APXs use electrons from ascorbate. The resulting dehydroascorbate is recycled through the ascorbate–glutathione cycle, which links H_2_O_2_ detoxification to the redox dynamics of the glutathione redox couple.

Glutathione is the most abundant low-molecular weight thiol-redox buffer in all eukaryotic organisms and most Gram-negative bacteria, including cyanobacteria and purple bacteria, and is present at millimolar concentrations (Fahey, 2001; Meyer *et al*., 2001). After synthesis in plastids and the cytosol (Pasternak *et al*., 2008; Maughan *et al*., 2010), reduced glutathione (GSH) is transported throughout the cell to fulfil a broad range of functions in metabolism and detoxification of xenobiotics and H_2_O_2_ (Meyer, 2008). Upon oxidation, two molecules of GSH convert to glutathione disulfide (GSSG). In the cytosol, mitochondria and plastids the glutathione redox potential (*E*_GSH_) is highly reduced with only nanomolar concentrations of GSSG present under non-stress conditions (Meyer *et al*., 2007; Schwarzländer *et al*., 2008). Such a high GSH/GSSG ratio is believed to be maintained by NADPH-dependent glutathione reductases (GRs). In contrast to bacteria, animals and yeast, plant genomes encode two GRs (Xu *et al*., 2013). In Arabidopsis, GR1 is present in the cytosol, the nucleus and peroxisomes (Reumann *et al*., 2007; Marty *et al*., 2009; Delorme-Hinoux *et al*., 2016). The second isoform, GR2, is dual-targeted to mitochondria and plastids (Creissen *et al*., 1995; Chew *et al*., 2003), as supported by proteomics analyses of purified organelles (Ito *et al*., 2006; Peltier *et al*., 2006). However, Yu and colleagues found no evidence for mitochondrial targeting of full-length GR2-YFP constructs and concluded exclusive plastidic localization (Yu *et al*., 2013). While we found that Arabidopsis mutants lacking cytosolic GR1 are fully viable (Marty *et al*., 2009), a deletion mutant lacking functional GR2 is embryo lethal (Tzafrir *et al*., 2004; Bryant *et al*., 2011). These observations raise important questions about the exact localization of GR2, the cause of lethality in *gr2* mutants, and about the mechanisms maintaining a highly negative glutathione redox potential (*E*_GSH_) in plastids and mitochondria.

Viability of Arabidopsis mutants lacking cytosolic GR1 is maintained by the NADPH-dependent TRX system (NTS) (Marty *et al*., 2009). In this case, electrons supplied by NADPH are transferred to GSSG via NADPH-dependent TRX reductase (NTR) and TRX. Arabidopsis contains two NTRs, NTRA and NTRB, which are both targeted to the cytosol, the nucleus and mitochondria (Reichheld *et al*., 2005; Marchal *et al*., 2014). In addition, plastids contain a remotely related bifunctional NTR, NTRC, which contains its own TRX domain (Serrato *et al*., 2004). NTRC has been shown to be responsible for transferring electrons from NADPH to 2-Cys PRX for H_2_O_2_ detoxification (Perez-Ruiz *et al*., 2017) and for redox regulation of ADP-glucose pyrophosphorylase (Michalska *et al*., 2009). While NTRC may be active in the dark and in heterotrophic tissues, classical non-fused plastidic TRXs are reduced by electrons from the photosynthetic electron transport chain via ferredoxin-dependent TRX reductase (FTR) which implies that they exercise their reductive capacity only in the light (Buchanan & Balmer, 2005). Whether and to what extent the different subcellular TRX systems contribute to organellar glutathione redox homeostasis is yet unknown. Beyond maintenance of a highly negative *E*_GSH_ through continuous reduction of GSSG, ATP-driven sequestration of GSSG to the vacuole by ATP-binding cassette (ABC)-transporters has been shown to provide an overflow valve for high cytosolic GSSG amounts in yeast (Morgan *et al*., 2013). Similarly, the ABC-transporter of the mitochondria Atm1 in yeast and its functional homologue ATM3 in Arabidopsis (systematic name ABCB25) transport GSSG, but not GSH, driven by ATP hydrolysis (Schaedler *et al*., 2014). The ATPase domain of Atm1 is orientated towards the mitochondrial matrix (Leighton & Schatz, 1995), indicating that these proteins export GSSG out of the mitochondria. In support of this, Arabidopsis *atm3* mutants were shown to have a more oxidized *E_GSH_* in the mitochondrial matrix compared to wild type (Schaedler et al 2014). However, the primary substrate of Atm1/ATM3 is thought to be GSH-bound persulfide for the biosynthesis of cytosolic iron-sulfur clusters (Kispal *et al*., 1999; Srinivasan *et al*., 2014). This raises the question of the ATM3’s relative contribution to other systems in GSSG clearance of the mitochondrial matrix.

Here, we address the role of GR2 in glutathione redox homeostasis in mitochondria and plastids. We localize GR2 by immunogold labelling and complement the lethal *gr2* null mutant in a compartment-specific manner. Those analyses provide clear evidence for dual localization of GR2 and for an essential role in plastids only. To address the question of why mitochondria are less affected than plastids by the lack of GR2, we focussed our attention on other possible mechanisms involved in the maintaining the *E*_GSH_ in the matrix. Using a series of physiological and genetic analyses we identify the involvement of alternative GSSG reduction systems and of GSSG export in *E*_GSH_ maintenance in the mitochondrial matrix.

## Materials and Methods

The following procedures are described in **Supporting Information Methods S1** and **S2**: Antibody production and gel blot analysis; Immunogold labelling and electron microscopy. Primers used in this work are given in **Supporting Information Table S1**.

### Plant material and growth conditions

The study was conducted with *Arabidopsis thaliana* ecotype Columbia-0 ([L.] Heynh.) as the wild-type (WT) control and the mutants *gr2-1* (SALK_040170, (Alonso *et al*., 2003), *emb2360-1* (Tzafrir *et al*., 2004), *ntra ntrb* (Reichheld *et al*., 2007), *rml1* (Vernoux *et al*., 2000) and *gsh1-1* (Cairns *et al*., 2006), *gr1-1* (Marty *et al*., 2009), and *atm3-4* (Bernard *et al*., 2009), all generated in the Col-0 background. Plants were grown under short day conditions: 8 h light at 22 °C and 16 h dark at 19 °C and light intensity was set to 120 µmol photons m^−2^ s^−1^. To induce flowering, plants were transferred to long day conditions: 16 h light at 22 °C and 8 h dark at 19 °C, light intensity was 160 µmol photons m^−2^ s^−1^.

To genotype WT and mutant plants, genomic DNA was extracted from the leaf tissue according to Edwards *et al*. (1991).

Seeds produced by a double heterozygous *gr2-1 rml1* plant were plated on phytagel as described before (Meyer & Fricker, 2000). For rescue experiments, the growth medium was supplemented with filter-sterilized GSH (Sigma-Aldrich) to a final concentration of 1 mM before gelling. Seedlings exhibiting a characteristic *rml1* phenotype after germination were transferred to GSH plates and further development was monitored for 12 d.

### Cloning and plant transformation

Standard molecular biology technologies like growth of bacteria, plasmid isolation and polymerase chain reaction (PCR) were applied according to (Sambrook *et al*., 1989). For cloning, all DNA fragments were amplified by PCR and blunt-end subcloned into pCAP (Roche Applied Science, Mannheim, Germany) with primers listed in Table S1. Accuracy of the cloned fragment was verified by sequencing by SeqLab (Göttingen, Germany). Different signal peptides were ligated into the pBinAR vector (Höfgen & Willmitzer, 1990) to achieve compartment-specific complementation. The transketolase targeting peptide (TK_TP_) sequence was cloned into pBinAR as described before (Wirtz & Hell, 2003). Similarly, the serine hydroxymethyltransferase target peptide (SHMT_TP_) was amplified and cloned behind the *35S* promoter of pBinAR using *Kpn*I and *Bam*HI. To verify targeting, both signal peptides were also fused to the N-terminus of roGFP. For compartment-specific complementation the GR2 sequence without its endogenous target peptide of 77 amino acids (aa) consistently determined by ChloroP 1.1 (http://www.cbs.dtu.dk/services/ChloroP/) and SignalP 3.0 (http://www.cbs.dtu.dk/services/SignalP/) (Bendtsen *et al*., 2004) was amplified from cDNA. Since the *gr2-1* mutants carried kanamycin resistance the constructs were transferred to the Basta^®^ resistance vector pBarA (a gift from Sabine Zachgo) using *Hind*III and *Eco*RI restriction sites. For constructs with the endogenous GR2 promoter, 2.1 kb were amplified from genomic DNA and used to replace the *35S* promoter in pBinAR with *Eco*RI and *Kpn*I.

Plant transformation was carried out by floral dip (Clough and Bent, 1998). For selection of positive transformants two-week-old plants were sprayed with Basta^®^ (200 mg l^−1^ glufosinate ammonium, Bayer Crop Science).

Arabidopsis plants were transformed with the vector pBinAR-SHMT-roGFP2-Grx1 as described earlier (Albrecht *et al*., 2014). For both WT and plastid-complemented *gr2* mutants, lines with incomplete mitochondrial targeting of roGFP2-Grx1 were selected.

### Protein purification and mitochondrial isolation

For recombinant protein expression, the pET28a and pETG10a constructs were transformed in *E. coli* HMS174 cells. Cells were grown at 37 °C to an OD600 nm of 0.8 in selective media containing 50 µg ml^−1^ kanamycin for pET28a or 100 µg ml^−1^ ampicillin for pETG10a. Protein purification was performed as reported before (Marty *et al*., 2009). Mitochondria were isolated from 13- to 16-day-old hydroponic Arabidopsis seedling cultures as described previously (Sweetlove *et al*., 2007).

### HPLC measurement of low-molecular weight thiols

Leaf material was harvested from six-week-old plants grown on soil under short-day conditions. Leaf tissue was snap frozen in liquid nitrogen, ground to fine powder and extracted. Extraction, derivatisation and quantification of low-molecular weight thiols by HPLC were done as described before (Meyer *et al*., 2007).

### Analysis of embryo development and pollen viability

Embryos, ovules or whole siliques were destained with Hoyeŕs solution (7.5 g gum arabic, 100 g chloral hydrate, 5 ml glycerol, 60 ml water) for at least 4 h. The embryos were analysed by differential interference contrast (DIC) microscopy with a Leica DM/RB microscope equipped with a DFC 320 camera (Leica Microsystems). Pollen from mature flowers were stained according to Alexander (1969) and analysed by bright field microscopy.

### Chlorophyll fluorescence measurement

Chlorophyll fluorescence was recorded after dark adaptation for at least 30 min with a pulse-amplitude modulated (PAM) fluorimeter (Junior PAM, Walz, Effeltrich, Germany). F_v_/F_m_ was calculated as a measure of the maximum potential quantum yield of photosystem II.

### Confocal laser scanning microscopy

Whole leaves of 3-week old Arabidopsis plants were placed in water on a slide and covered with a coverslip. Leaves were imaged on Zeiss confocal microscopes LSM510META or LSM780 (Carl Zeiss MicroImaging, Jena, Germany) equipped with lasers for 405- and 488-nm excitation. Images were collected with either a 25x lens (Plan-Neofluar 25x/0.8 Imm corr, Zeiss) or a 63x lens (C Apochromat 63x/1.2 W corr, Zeiss). For localization studies GFP was excited at 488 nm and emission was collected with a 505-530 nm band-pass filter. Chlorophyll autofluorescence was excited at 405 nm and recorded above 630 nm. For counterstaining mitochondria leaf tissue was incubated with 1 µM tetramethylrhodamine methyl ester (TMRM) for 30-60 min. TMRM was excited at 543 nm and recorded at 582-646 nm. For glutathione labelling in roots of Arabidopsis seedlings intact seedlings were incubated with 100 µM monochlorobimane (MCB) and 50 µM propidium iodide (PI) for 30 min and imaged as reported earlier (Meyer *et al*., 2001). For ratiometric imaging of *E*_GSH_ roGFP2-Grx1 was excited in multi-track mode with line switching between 488 nm illumination and 405 nm illumination. The roGFP2 fluorescence was collected with a 505–550 nm emission band-pass filter. Ratiometric image analysis was done as reported previously (Schwarzländer *et al*., 2008) using a custom Matlab programme (Fricker, 2016).

## Results

### *gr2* T-DNA insertion allele causes early embryonic lethality

To investigate the role of GR2 for glutathione redox homeostasis, T-DNA insertion lines for *GR2* were identified and characterized. The T-DNA insertion lines *emb2360-1*, *emb2360-2*, *emb2360-3* (Tzafrir *et al*., 2004), and *gr2-1* (SALK_040170) were obtained from NASC. The T-DNA insertion in *gr2-1* was localized in the 8^th^ intron (Fig. 1a) and the insertion site was confirmed by sequencing using a T-DNA left border primer. A population of 133 plants derived from a heterozygous *gr2-1* plant segregated with 85 kanamycin-resistant (Kan^R^) : 48 kanamycin-sensitive plants. This segregation of the antibiotic resistance in a 2:1 ratio (χ^2^ = 0.45, *P* = 0.79) indicates that homozygous *gr2-1* segregates as a single locus and is sporophytic lethal. Genotyping of soil-grown Kan^R^ plants confirmed that all Kan^R^ plants were heterozygous *gr2-1*, demonstrating that Kan^R^ is caused by the T-DNA in the *GR2* locus. Microscopic examination of immature siliques of heterozygous *gr2-1* mutants showed that 25 % (31:97, χ^2^ = 0.04 for a 1:3 segregation, *P* = 0.98) seeds were white with embryos arrested at globular stage of development (Fig. 1d,e). Consistent segregation patterns and phenotypes were found for the three *emb2360* alleles (http://seedgenes.org/SeedGeneProfile_geneSymbol_EMB_2360.html), providing independent evidence for a link between GR2 function and embryonic lethality. All further experiments were done with *gr2-1*, which is from now referred to as *gr2*.

**Fig. 1.**
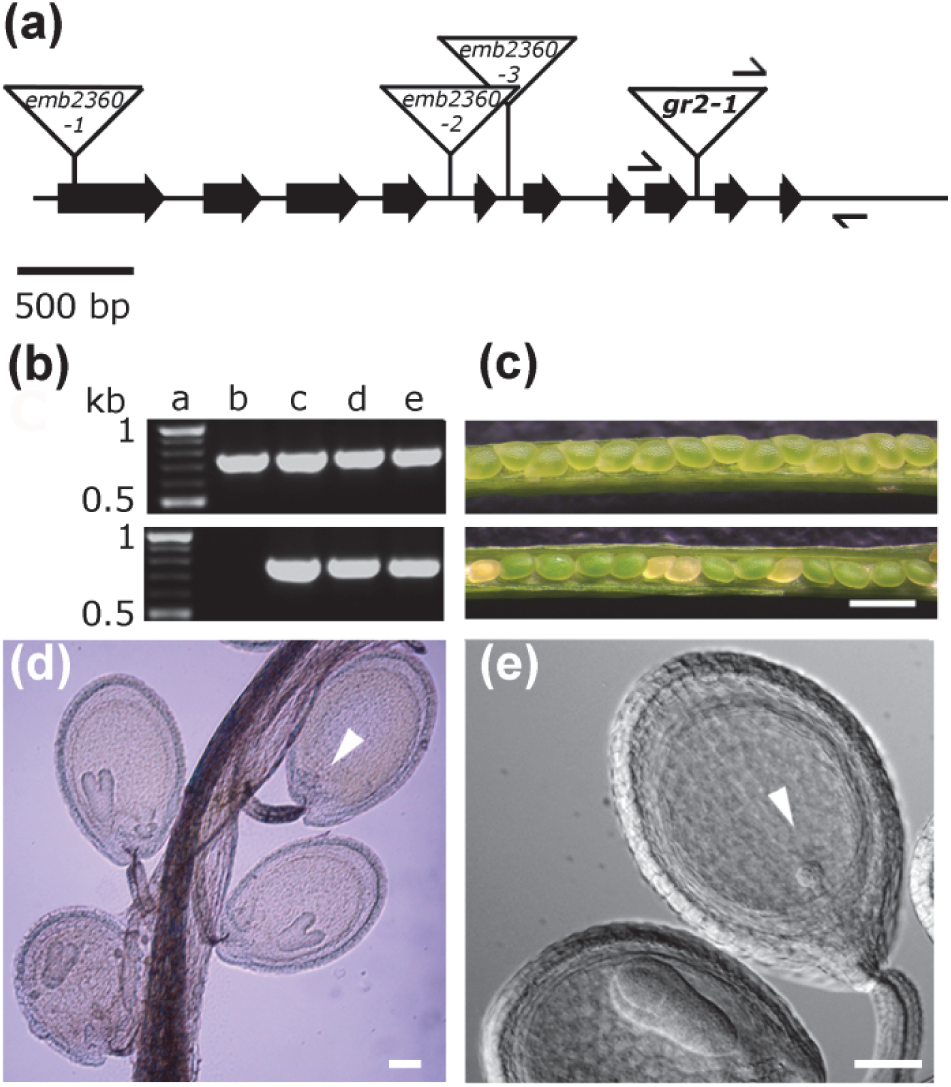
Isolation and phenotypic characterization of *gr2* null mutants. (a) Exon/intron structure of the Arabidopsis *GR2* gene (At3g54660). Exons are represented by large arrows and the triangles show the T-DNA insertions for three *emb2360* alleles (http://seedgenes.org/SeedGeneProfile_geneSymbol_EMB_2360.html) and the allele *gr2-1*. Primer binding sites for genotyping of *gr2-1* are indicated by black arrows. (b) Genotype analysis of different *gr2-1* lines. Upper panel: PCR performed with *GR2* gene specific primers. Lower panel: PCR with a gene-specific primer and a primer for the left T-DNA border. Lane a: DNA marker: size is indicated in kb; Lane b: WT; Lanes c-e: heterozygous *gr2-1* mutants. (c) Embryo-lethal phenotype associated with *gr2-1*. Immature siliques from self-fertilized WT (upper panel) and heterozygous *gr2-1* mutants (lower panel) were opened to observe segregation of seed phenotypes. Bar, 500 µm. 25 % of the seeds in the *gr2* silique show a lethal phenotype visible as white seeds. (d-e) Developing seeds of heterozygous *gr2-1* mutants were cleared using Hoyer’s solution and analysed by bright field microscopy (d) and DIC microscopy (e). Arrows indicate white ovules containing embryos arrested at globular stage. Bars, 50 µm.

### GR2 solely targeted to plastids rescues the lethal *gr2* phenotype

With the precedence of contradicting results regarding the subcellular localization of GR2 (Chew *et al*., 2003; Yu *et al*., 2013) we attempted an orthogonal method to explore localization and raised polyclonal antibodies against GR2 (Fig. S2b). Immunogold labelling of GR2 in leaf tissue of WT plants independently demonstrates its dual localization (Fig. 2a,d). The early embryonic lethal phenotype of *gr2* may thus be caused by mitochondrial or plastidic defects, or both. To resolve this, *gr2* mutants were complemented with organelle-specific *GR2* constructs. The endogenous signal peptide of 77 aa was replaced by the signal peptides SHMT for mitochondria or TK_TP_ for plastids (Schwarzländer *et al*., 2008; Albrecht *et al*., 2014) (Fig. S1).

**Fig. 2.**
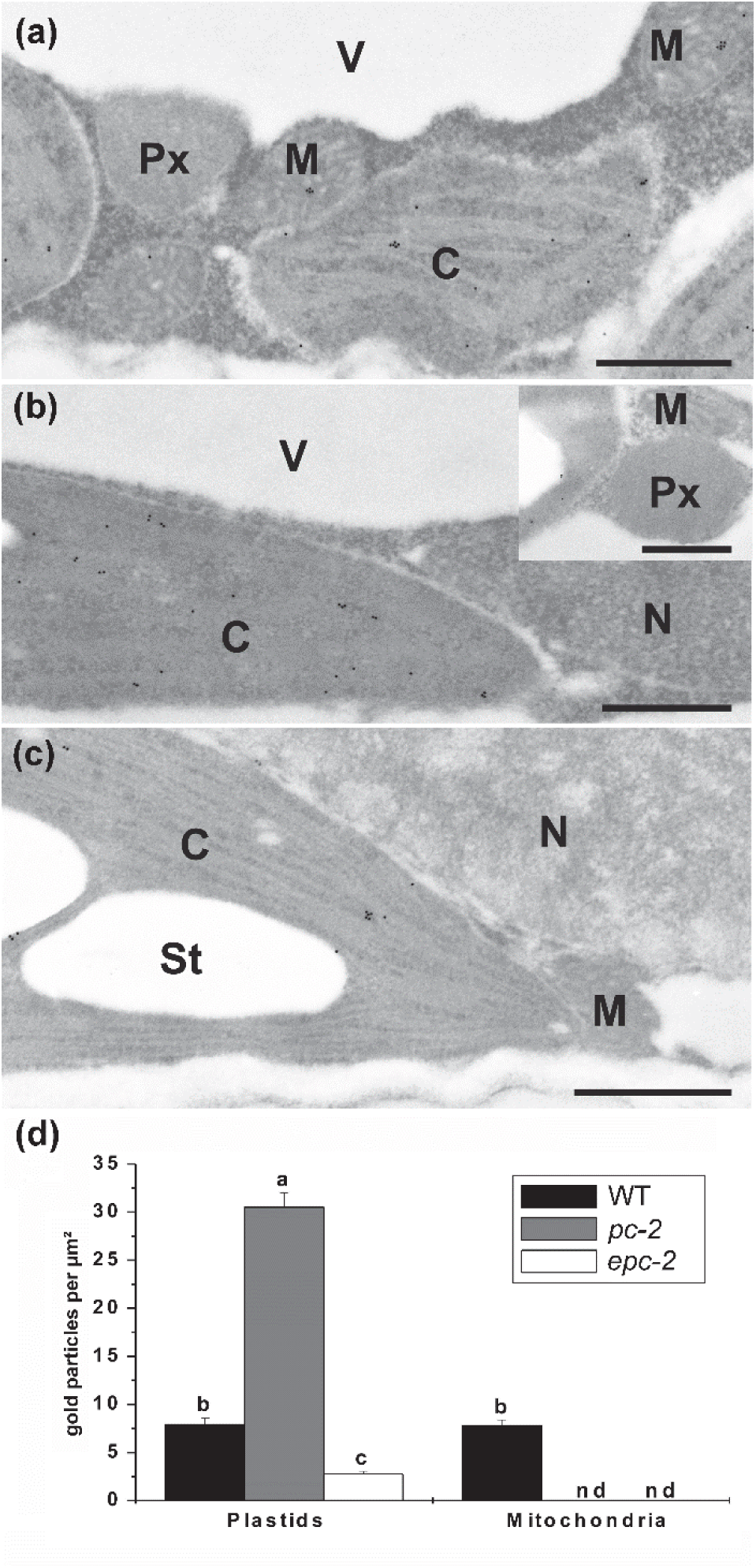
Immunogold localization of GR2 in plastid complemented *gr2* deletion lines. Fixed and dissected mature leaves were probed with a primary antibody raised against Arabidopsis GR2. C: chloroplast; M: mitochondrion: Px: peroxisome; V: vacuole; N: nucleus; St: starch grain. Bars, 1 µm. (a) WT. (b) Line *pc-2* (‘plastid complemented’) with TK_TP_-Δ_1-77_GR2 expressed from the *35S* promoter. (c) Line *epc-2* with TK_TP_-Δ_1-77_GR2 expressed from the endogenous GR2 promoter. (d) Quantitative analysis of gold particles observed in electron microscopy micrographs. Values are means ± SE and document the amount of gold particles per µm^2^ in the respective organelle. Data were analysed by the Kruskal-Wallis test, followed by post hoc comparison according to Conover. Different lowercase letters indicate significant differences (*P* < 0.05). *n* > 60. nd = not detected.

Compartment-specific complementation of *gr2* was initially done with GR2 constructs driven by the *CaMV 35S* promoter (*35S_pro_*) (Fig. S1). T2 transformants (named *pc* for <u>p</u>lastid <u>c</u>omplementation) were selected with Basta® and genotyped for the *gr2* locus to screen for successful complementation of the lethal phenotype. Transformation with *35S_pro_:TK_TP_-Δ_1-77_GR2* resulted in 22 % of the progeny being homozygous for *gr2* consistent with the theoretical value of 25 % rescued homozygous mutants (Table S2). From these complemented mutants three independent lines, named *pc-1, pc-2, and pc-3*, were selected and characterized in more detail (Fig. S2). Protein gel blots of WT leaf extracts consistently revealed two distinct protein bands of ∼53 and ∼110 kDa (Fig. S2b). The 110 kDa protein band was less intense than the 53 kDa band and most likely represents a GR2 homodimer. The unprocessed WT GR2 protein including the endogenous signal peptide of 77 aa has a predicted size of 61 kDa. The 53 kDa band detected in the protein gel blot likely corresponds to the mature processed protein with a predicted size of 52.7 kDa. This result indicates that cleavage of the endogenous GR2 signal peptide is the same for plastids and mitochondria. In *pc* lines over-expressing TK_TP_-GR2, a very strong increase in both bands was observed without significant change in the relative distribution between the two bands. Moreover, in all *35S_pro_:GR2* over-expression lines several additional protein bands with masses below that of GR2 may indicate partial degradation of GR2. Total GR activity in leaf tissue extracts was 49 nmol min^−1^ mg^−1^ for WT and about 700 nmol min^−1^ mg^−1^ for the *pc* lines (Fig. S2c). This 15-fold increase is consistent with the pronounced increase in GR2 abundance observed in the *pc* lines. All *pc* lines showed WT-like GSH and GSSG levels with no changes in the GSH/GSSG ratio detectable by HPLC-based thiol analysis (Fig. S2d). At random time points in their development several T1 plants complemented with *35S_pro_:TK_TP_-Δ_1-77_GR2* unexpectedly showed wilted, dwarfed and purple coloured phenotypes, ultimately resulting in plant death before seed setting (Fig. S2a). This phenotype occurred irrespective of whether the genetic background of the respective plants was WT, *gr2*^+/−^ or *gr2*^−/−^. The occurrence of plant death in WT and *gr2*^+/−^ plants strongly suggests that the phenotype was caused by co-suppression of the endogenous *GR2* gene together with the transgene.

To avoid potential artefacts resulting from extremely high levels of GR2 protein and also to minimize the risk of silencing, *gr2* mutants were complemented with *GR2* driven by its endogenous promoter (Fig. S1). In this case, however, only 10 % of the progeny, named *epc*, were found to be homozygous for *gr2* (Table S2). The lower frequency of the complementation with GR2 driven by the endogenous promoter is most likely due to the much lower activity of this promoter compared to the *35S_pro_*. In some cases, the activity of the used *GR2_pro_*may not be sufficient to achieve full complementation and thus cause premature abortion of homozygous *gr2* embryos.

In contrast, none of the transgenic lines transformed with either *35S_pro_:SHMT_TP_-Δ_1-77_GR2* or *GR2_pro_:SHMT_TP_-Δ_1-77_GR2* for mitochondrial targeting of GR2 was found to be a homozygous knockout for *gr2* (Table S2). Thus, only *TK_TP_-Δ_1-77_GR2* constructs controlled by either the endogenous *GR2_pro_* or *35S_pro_* rescued the lethal *gr2* mutant, indicating that only plastid-localized GR2, but not the mitochondria-localized GR2, is essential for development. Correct targeting of both GR2 plastid complementation constructs was confirmed by immunogold labelling (Fig. 2b-d).

Three independent homozygous *gr2* mutants complemented with *GR2_pro_:TK_TP_-Δ_1-77_GR2* constructs were isolated and named *epc-1, epc-2*, and *epc-3*, respectively. When grown on soil under short day conditions, all three *epc* lines showed WT-like phenotypes in growth and development (Fig. 3a). GR2 protein levels in T1 plants with different genotypes with respect to the *gr2* locus were determined by protein gel blot analysis (Fig. 3b). Similar to WT and the *35S_pro_* complemented lines, protein gel blot analysis of total protein extracts consistently revealed two bands of ∼110 kDa and ∼53 kDa, respectively, with the 53 kDa band being much more intense. In addition to the 53 kDa and 110 kDa bands, protein extracts of *TK_TP_-Δ_1-77_GR2* transformants showed a third band with a size of ∼55 kDa. The appearance of this band suggests that import processing of the TK_TP_-Δ_1-77_GR2 protein differed from WT GR2. Indeed, ChloroP predicts TK_TP_-Δ_1-77_GR2 processing to result in a mature protein with a mass of 54.2 kDa. In all three homozygous *gr2* complementation lines only the 55 kDa protein was identified while the 53 kDa band was absent, further confirming at biochemical level that all three selected *epc* lines were indeed *gr2* null mutants successfully complemented with TK_TP_-Δ_1-77_GR2. Furthermore, the absence of a larger band of 61 kDa indicates that plastid import of GR2 was highly efficient without detectable traces of non-processed cytosolic protein. Protein gel blot analysis of plants transformed with *GR2_pro_:SHMT_TP_-Δ_1-77_GR2* showed the same two-band pattern found in WT (Fig. 3b, lane d). Lack of additional bands suggests that the processed WT GR2 and processed SHMT_TP_-Δ_1-77_GR2 proteins have similar sizes, which is also supported by bioinformatics prediction by SignalP 3.0.

**Fig. 3.**
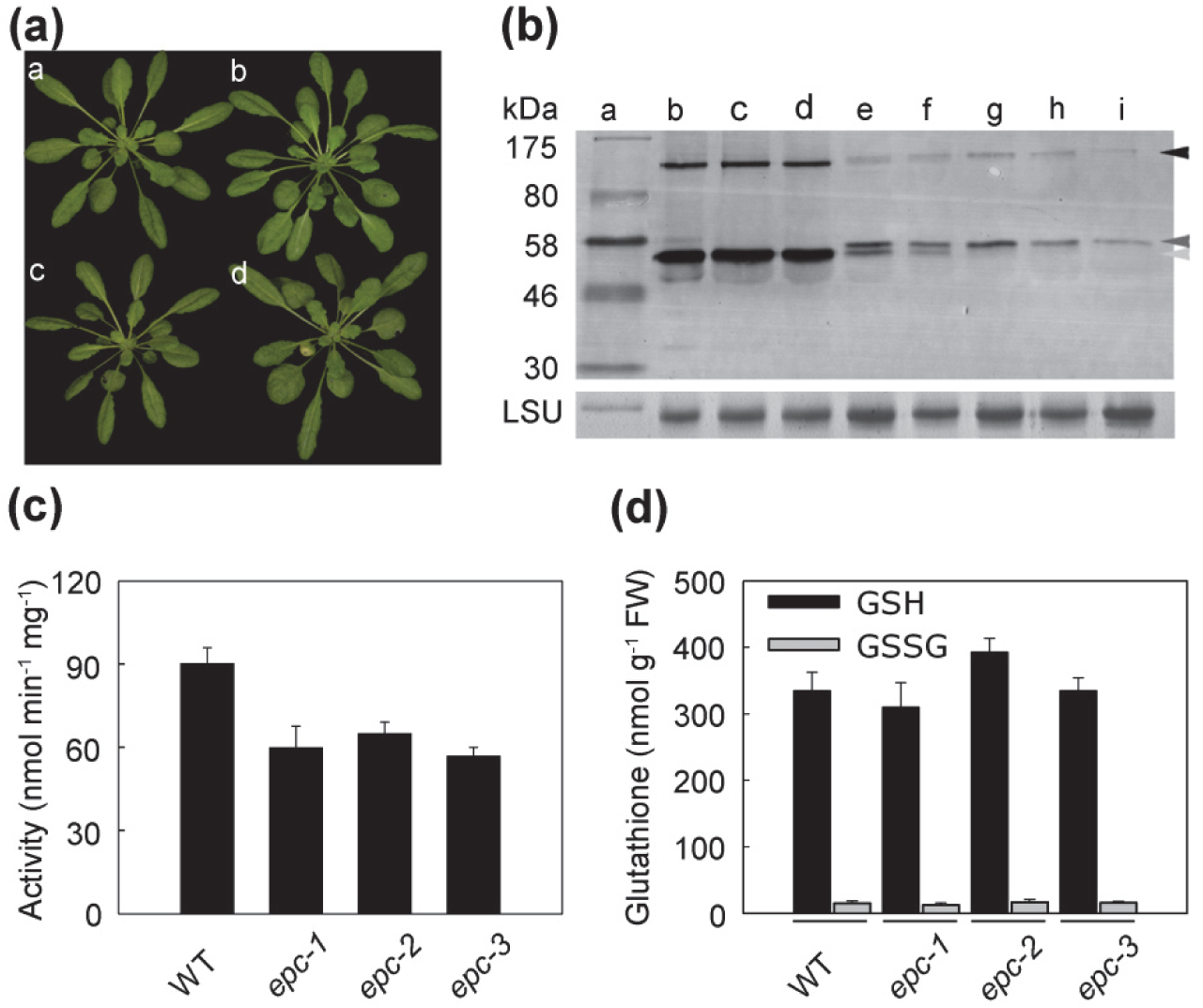
Characterization of lines expressing plastid-targeted GR2 controlled by its endogenous promoter. (a) Growth phenotypes of WT (a) and three independent homozygous *gr2* deletion mutants complemented with *GR2_pro_:TK_TP_-Δ_1-77_GR2*: *epc-1* (b), *epc-2* (c), *epc-3* (d). All transgenic plants were from the Basta^®^-selected T1 generation. (b) Protein gel blot analysis of transgenic Basta^®^-selected T1 lines transformed with complementation constructs driven by the endogenous *GR2* promoter and targeted to either mitochondria or plastids. Loading was as follows: a: Pre-stained molecular mass standard, b: WT, c: WT transformed with *GR2_pro_:SHMT_TP_-Δ_1-77_GR2*, d: *gr2*^+/−^ transformed with *SHMT_TP_-Δ_1-77_GR2*, e: WT transformed with *GR2_pro_:TK_TP_-Δ_1-77_GR2*, f: *gr2*^+/−^ transformed with *GR2_pro_:TK_TP_-Δ_1-77_GR2*, g-i: three independent homozygous *gr2* deletion lines transformed with *GR2_pro_:TK_TP_-Δ_1-77_GR2* (*epc-1*, *epc-2* and *epc-3*). Arrows indicate protein bands with the size of 110 kDa (black), 55 kDa (dark grey) and 53 kDa (light grey). (c) GR activity in total protein extract from leaves. Proteins were extracted from WT and T2 plants of the homozygous *gr2* lines *epc-1*, *epc-2* and *epc-3*. Means ± SD of three independent plants of each line are shown. (d) Contents of oxidized and reduced glutathione. After extraction of leaf tissue of 5-week-old plants reduced glutathione (GSH) and glutathione disulfide (GSSG) were determined by HPLC. Means ± SD of 6 independent plants of each line are shown.

### Minute amounts of functional GR2 are sufficient for growth

GR activity of total protein extracts of the *epc* lines was 60 nmol min^−1^ mg^−1^ and thus ∼30 % lower than WT GR activity of 90 nmol min^−1^ mg^−1^ (Fig. 3c). Both, protein gel blot analysis and GR activity measurements of all three independent *epc* lines, showed that the amount of GR2 protein was significantly lower than in WT. This indicates that either promoter and/or enhancer elements other than the 2.1 kb fragment used here, may contribute to control the GR2 expression in WT plants, or that positional effects may have led to partial repression of GR2 in *epc* lines. The content of GSH and GSSG was comparable to WT plants (Fig. 3d).

The GR activity in *epc* lines is due to the *GR2* transgene and the endogenous GR1 activity, which had been reported to account for 40 to 60 % of the total GR activity in leaves (Marty *et al*., 2009; Mhamdi *et al*., 2010). To further separate these activities and to better assess the requirement for GR2 necessary for normal growth, we generated a double heterozygous *gr1^+/−^ gr2*^+/−^ plant and complemented this with the same *GR2_pro_:TK_TP_-Δ_1-77_GR2* construct used before. From the F2 progeny we selected two homozygous *gr1 gr2* double knockouts that expressed GR2 exclusively in the plastids (lines *epc-9* and *epc-27*). These *gr1 gr2* plants did not show any obvious phenotype in their growth and development during the vegetative phase under standard growth conditions (Fig. 4a). The total GR activity in leaf extracts was only between 10 ± 2.6 and 1.5 ± 0.1 nmol mg^−1^ min^−1^ compared to 53.2 ± 3.5 nmol mg^−1^ min^−1^ in WT leaves (Fig. 4b). For comparison, *gr1* mutants and the plastid-complemented *gr2* mutant (*epc-2*) showed intermediate GR-activities of ± 2.6 and 24.2 ± 2.6 nmol mg^−1^ min^−1^, respectively (Fig. 4b). Lack of GR1 in *gr1-1* and in *gr1 gr2* double deletion mutants consistently resulted in an increase in total glutathione (Fig. 4c). Further analysis of the glutathione pool revealed that this increase was largely due to an increase in GSSG, which was about six-fold higher compared to WT controls (Fig. 4d). If at all, plastid-specific complementation of *gr2* resulted in only a minor increase in glutathione (Figs. 3d and 4c). To further test for mitochondrial GR activity, we first isolated mitochondria from the plastid complemented line *epc-2* and determined the specific GR activity in mitochondrial extracts. While mitochondria isolated from WT plants had a specific GR activity of 47.9 nmol min^−1^ mg^−1^, mitochondria from *epc-2* contained an activity of 2.4 nmol min^−1^ mg^−1^ (Fig. 5a). Identity of mitochondrial preparations was confirmed by detection of the mitochondrial marker peroxiredoxin II F (PRXII F; (Finkemeier *et al*., 2005)). Further analysis of the respective extracts indicated that GR2 was absent in *epc-2* mitochondria (Fig. 5c). Hybridization of the respective protein blots with a GR1 antibody (Marty *et al*., 2009) revealed that the mitochondrial fraction also contained GR1 (Fig. 5c) which can be accounted for by the presence of peroxisomes that contain GR1, in mitochondrial preparations from Arabidopsis seedlings (Sweetlove *et al*., 2007). No GR activity was found in mitochondria isolated from line *epc-27* (Fig. 5b). Protein gel blots confirmed absence of both GR1 and GR2 from *epc-27* mitochondria (Fig. 5d).

**Fig. 4.**
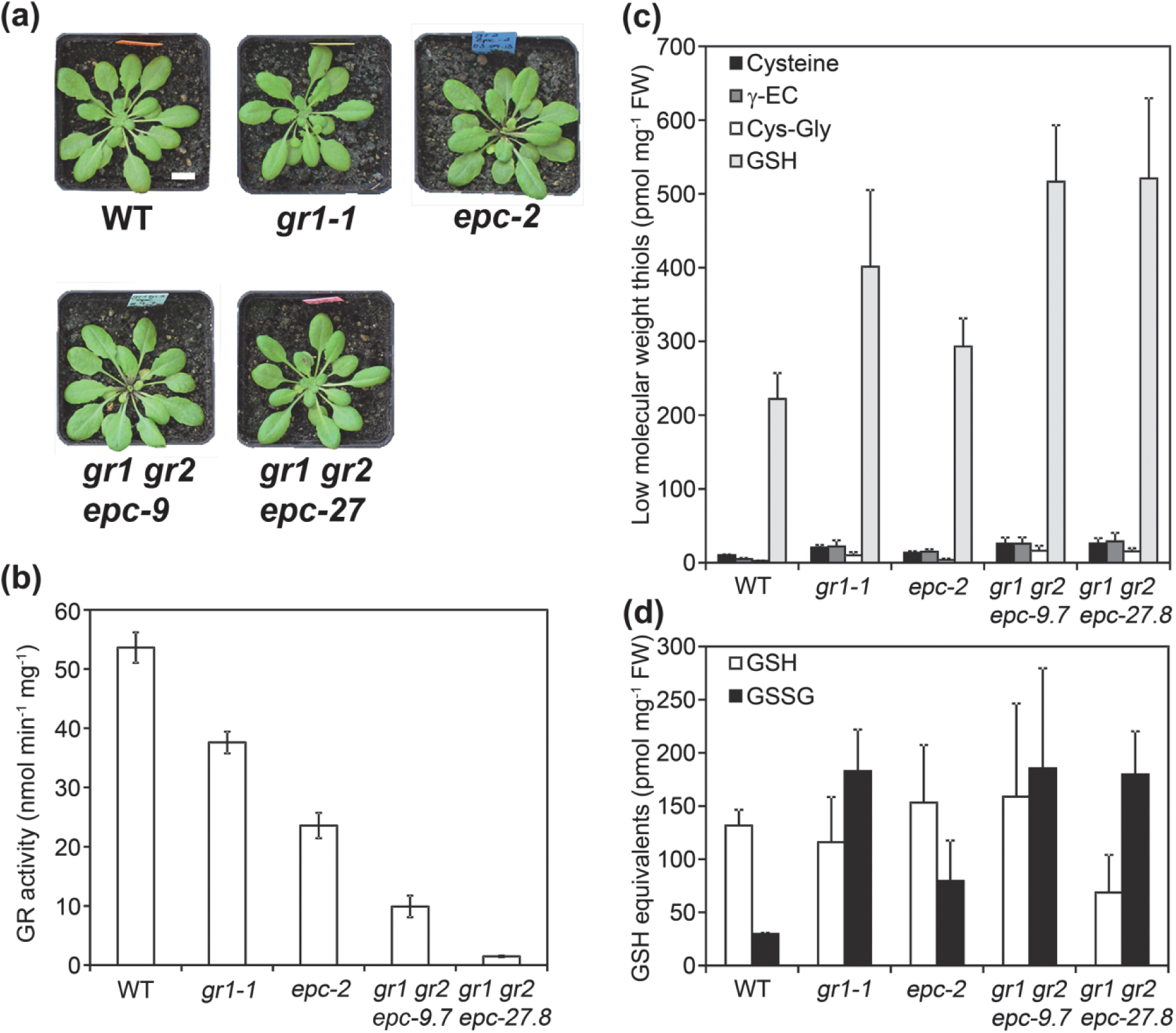
Glutathione reductase activity and low molecular weight thiols of *gr1 gr2* double homozygous mutants complemented with plastid-targeted GR2. (a) Rosette phenotypes of plants grown on soil for six weeks under short day conditions (8 h : 16 h, light : dark). Bar, 1 cm. (b) GR activity in total protein extracts from leaves. Proteins were extracted from WT, *gr1*, plastid complemented *gr2* (*epc-2*) and two *gr1 gr2* double mutants (*epc-9* and *epc-27*) that were complemented with *GR2_pro_:TK_TP_-Δ_1-77_GR2*. Means ± SD of three independent plants of each complemented line are shown. (c) Low-molecular weight thiols analysed by HPLC from leaf extracts. GSH in this case refers to total glutathione. Means ± SD, *n* = 3. (d) Reduced glutathione (GSH) and glutathione disulfide (GSSG) determined by HPLC in leaf extracts. Means ± SD, *n* = 3.

**Fig. 5.**
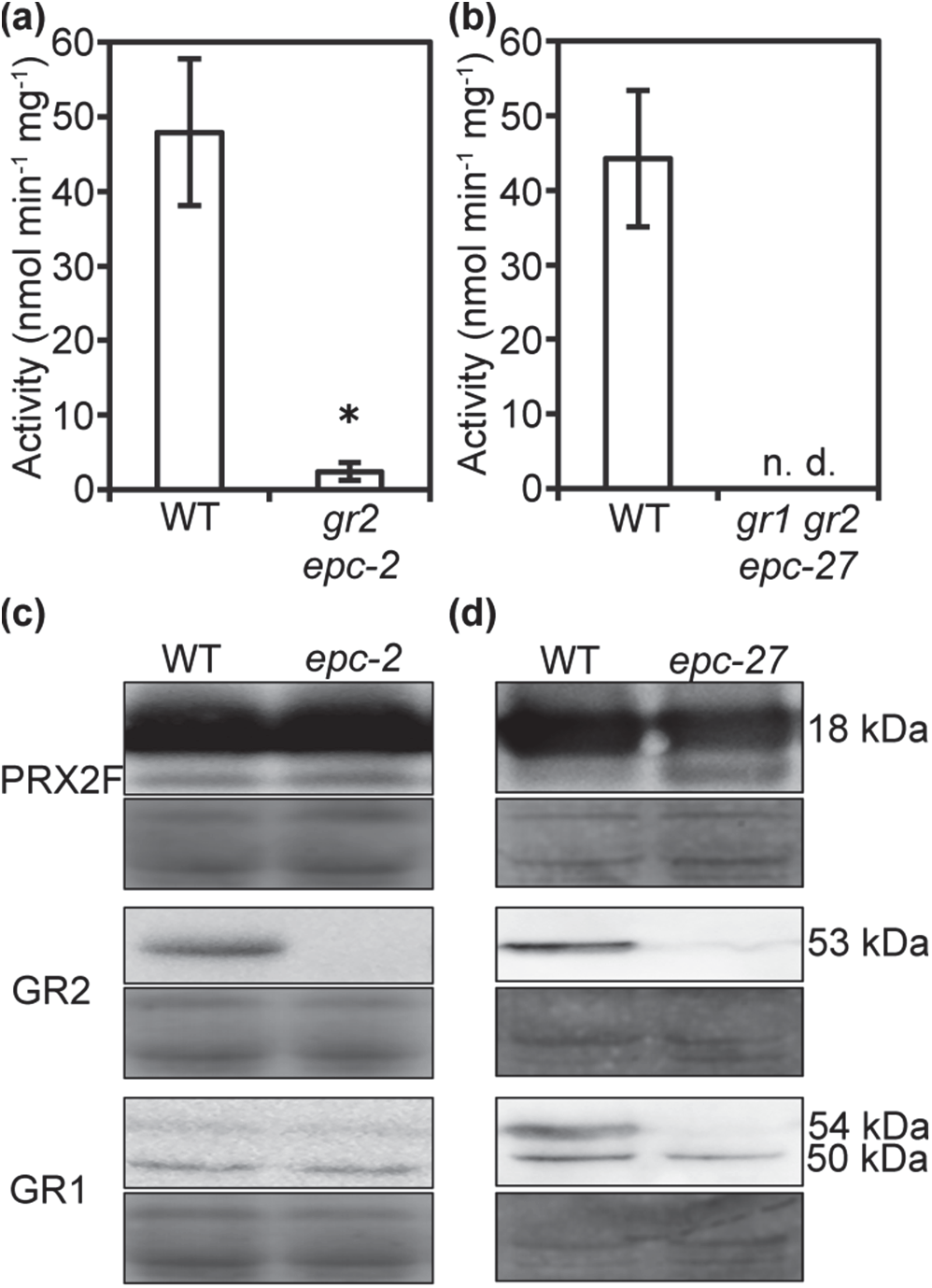
Glutathione reductase activity in isolated mitochondria. Intact mitochondria were isolated from two-week old hydroponically grown Arabidopsis plants (WT and *epc-2* lacking endogenous GR2, or *epc-27* lacking both endogenous GRs, respectively). Activities represent the means from four independent preparations with six technical replicates. Error bars represent SD. (a, b) GR activity. n.d. = not detected; * indicates *P* < 5*10^−4^. (c, d) Protein gel blots of PRXII F, GR2 and GR1 in mitochondrial preparations. Loading controls were stained with amido black. In addition to GR1 (54 kDa), the GR1 antibody detected also a smaller protein of about 50 kDa (Fig. 5d). This band, however, appears to result from a side activity against another mitochondrial protein that is detectable in concentrated isolated mitochondria, but not in whole leaf extracts (Fig. S3b). Mitochondria contain two lipoamide dehydrogenases (MTLPD1 and MTLPD2) that are closely related to GR and are both predicted with a molecular mass of 49.9 kDa after cleavage of the mitochondrial target peptide (Lutziger & Oliver, 2001). Thus, it is likely that the additional band results from a cross reaction of the antibody with MTLPDs.

### Glutathione deficiency partially suppresses embryo lethality of *gr2*

Early embryonic lethality of *gr2* null mutants may be explained by lack of appropriate backup systems for GSSG reduction, or an additional, yet unknown, moonlighting function (Jeffery, 2009) of plastidic GR2. To elucidate whether the early embryonic lethal phenotype is caused solely by accumulation of GSSG in plastids, we reasoned that GSH-deficiency would necessarily go along with deficiency of GSSG and thus might suppress the early embryonic lethal phenotype of *gr2*. To test this hypothesis, *gr2* was crossed with the two GSH-deficient mutants *rml1* and *gsh1-1*, which are both compromised in the first step of GSH biosynthesis catalysed by glutamate-cysteine ligase (GSH1) and contain strongly decreased glutathione levels (Vernoux *et al*., 2000; Cairns *et al*., 2006). While *rml1* completes embryogenesis and germinates, the null mutant *gsh1-1* reaches U-turn stage before it dies (Vernoux *et al*., 2000; Cairns *et al*., 2006). While no double homozygous *rml1 gr2* could be found (Fig. S4) the *gr2 gsh1* cross and selfing of a double heterozygous plant resulted in approximately 24 % aborted seeds albeit with a distinct phenotypic variation (Fig. 6c). Closer inspection of the aborted seeds revealed that about 1/4 of seeds were aborted later than the typical *gr2* embryo, which were seen as white turgescent ovules (Fig. 6c). Indeed developing seeds collected from a selfed double heterozygous *gr2*^+/−^ *gsh1*^+/−^ plant showed a segregation of their embryos in 334 U-turn : 92 globular : 30 torpedo (12:3:1, χ^2^ = 0.76, *P* = 0.34) (Fig. 6d-g). This result indicates that GSH-deficiency caused by *gsh1* partially suppressed the early embryo-lethal *gr2* phenotype extending development until the torpedo stage. This points to GSSG toxicity, as opposed to an oxidative shift in *E*_GSH_, as a cause of the embryo lethality in the absence of GR2 from the plastid stroma.

**Fig. 6.**
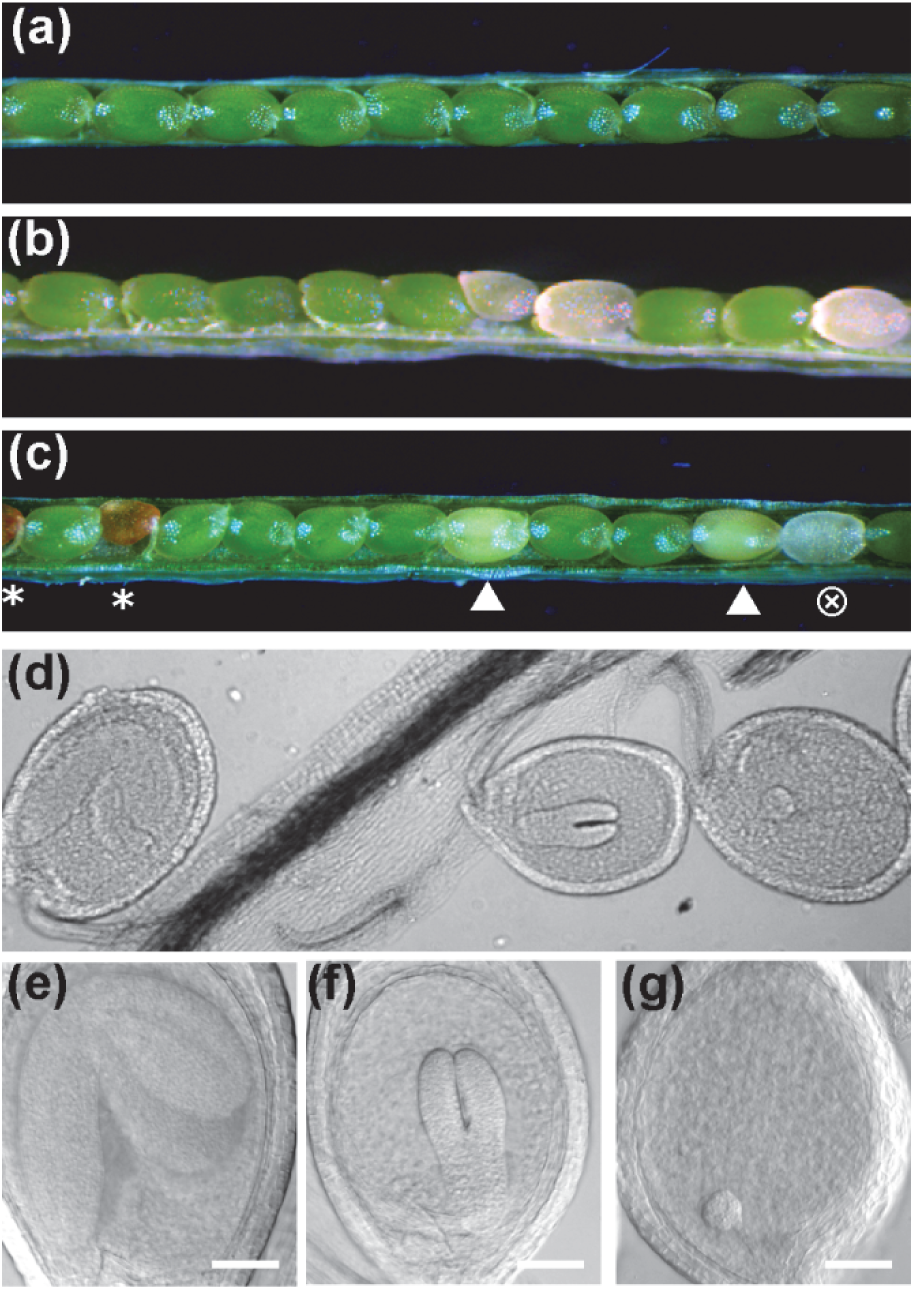
Characterization of *gr2 gsh1* double mutants. (a) WT silique at 14 d after fertilization (DAF). (b) Silique of self-fertilized heterozygous *gr2* plant displaying 25 % aborted seeds. (c) Silique of self-fertilized double heterozygous *gr2*^+/−^ *gsh1*^+/−^ plant displaying segregation of green WT seeds, partially bleached *gsh1* seeds (▲), brownish early aborted *gr2* seeds (*), and transparent seeds (⊗) that remain fully turgescent significantly longer than *gr2* seeds. (d-g) DIC images of ovules developed in a silique of a self-fertilized double heterozygous *gr2*^+/−^ *gsh1*^+/−^ plant 14 DAF. Bars, 50µm.

### The mitochondrial glutathione pool of plastid-complemented *gr2* mutants shows increased oxidation

Full viability of plastid complemented *gr2* plants suggests that mitochondria can maintain their function in the absence of GR2. This suggests that they are either able to cope with a fully oxidized glutathione pool made up of GSSG, or other mechanisms of maintaining reduction exist. Candidate mechanisms include GSSG export from the matrix for reduction in the cytosol as well as an alternative enzymatic system for efficient internal GSSG reduction. To address this question, we tested whether *E_GSH_* in the mitochondrial matrix of mutants lacking GR2 in mitochondria is more oxidized than in WT controls. Plastid complemented *gr2*^−/−^ mutants (*epc-2*) were transformed with the *E*_GSH_ biosensor roGFP2-Grx1 targeted to the mitochondrial matrix (Albrecht *et al*., 2014). roGFP2 constructs were expressed from *35S_pro_* and we deliberately selected strongly expressing lines in which part of the sensor proteins remained in the cytosol and the nucleus (Fig. 7 and Fig. S5). The resulting dual localization of roGFP2-Grx1 allowed simultaneous ratiometric imaging of roGFP2 in both compartments by confocal microscopy. Because the roGFP2-Grx1 in the cytosol and the nucleus provides an internal control for close-to-complete sensor reduction (Meyer *et al*., 2007; Schwarzländer *et al*., 2008) the ratiometric images are informative without any further treatment with reducing and oxidizing compounds. While the merge of the raw images collected after excitation at 405 nm and 488 nm in WT seedlings show similar green colour in cytosol and mitochondria (Fig. 7a) the same merge for the plastid complemented *gr2*^−/−^ mutant shows a pronounced difference: Cytosol and nucleus appear in green like in the WT control, while the mitochondria appear in yellow because the fluorescence in the 488 nm channel decreased and the fluorescence in the 405 nm channel increased (Fig. 7b). The differential behaviour of the signals in mitochondria and the cytosol is even more obvious after ratiometric analysis, which shows a partial oxidation of roGFP2-Grx1 in the GR2-deficient mitochondria. The degree of sensor oxidation can be estimated to approximately the midpoint of the sensor, i.e. around 50 %. This corresponds to a pronounced shift in mitochondrial matrix *E*_GSH_ of about 30 mV, but is remains far from a complete oxidation of the matrix glutathione pool.

**Fig. 7.**
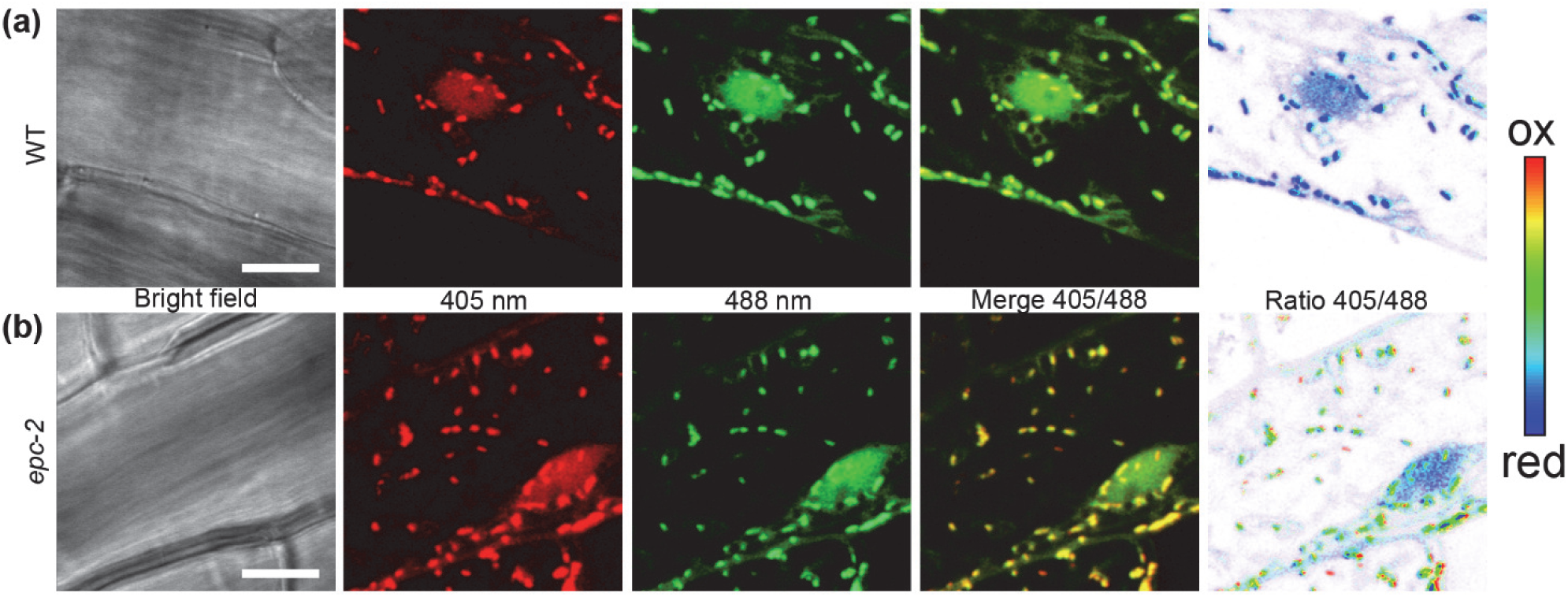
Simultaneous imaging of the local glutathione redox status in mitochondria and the cytosol. The ratiometric redox probe roGFP2-Grx1 was expressed with a SHMT_TP_ mitochondrial targeting sequence driven by a *35S_pro_*. After transformation lines with residual roGFP2-Grx1 in the cytosol were selected and used for simultaneous redox imaging in mitochondria and the cytosol. Images show hypocotyl cells in WT seedlings (a) or plastid complemented *gr2*^−/−^ seedlings (b). Images taken after excitation at 405 nm are false colour coded in red and images taken after excitation at 488 nm are coded in green in order to visualize a pronounced colour change in the merge image from green to yellow in the mitochondria. Bars, 10 µm.

### The mitochondrial ABC-transporter ATM3 has a critical function in GR2-deficient mitochondria

Export of GSSG from the mitochondrial matrix and subsequent reduction by cytosolic GR1 activity would provide an alternative mechanism to adjust the matrix E_GSH._ To elucidate whether the GSSG export ability of ATM3 could help to maintain low GSSG levels and support the mitochondrial GR2 activity, we crossed the mutant *gr2 epc-2* deficient in mitochondrial GR2 with *atm3-4*, and isolated the double mutant. *atm3-4* is a relatively weak mutant allele, with low expression of functional *ATM3* due to a 39-nucleotide deletion in the promoter (Bernard *et al*., 2009). The *atm3-4 gr2 epc2* double mutant showed an enhanced phenotype compared to both the *gr2 epc2* and *atm3-4* parents, with short roots at seedling stage and smaller rosettes when grown on soil (Fig. 8). Leaves appeared chlorotic and displayed a significantly lower F_v_/F_m_ (Fig. 8d). These observations suggest synergistic action of the two mutations with lack of mitochondrial GR2 causing increased GSSG levels in the matrix that cannot be efficiently decreased when ATM3 protein levels are strongly depleted.

**Fig. 8.**
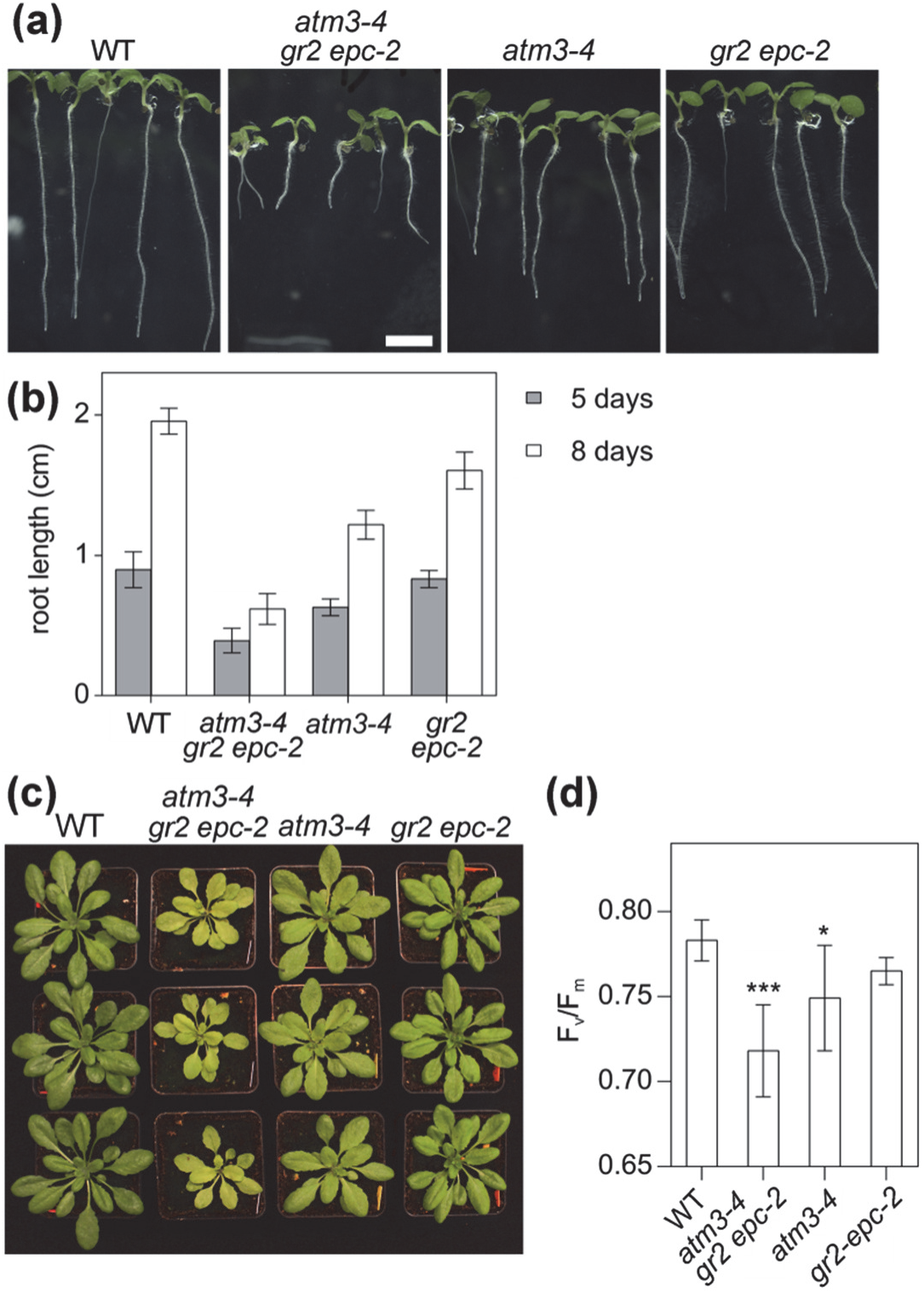
*atm3-4 gr2 epc2* double mutants show a semi-dwarf and chlorotic phenotype. (a) 8-d-old seedlings of the double mutant compared to WT and single mutants. Bar, 0.5 cm. (b) Root length of the mutants compared to WT. Plants were grown on ½ MS medium containing 0.5 % sucrose and 0.8 % phytagel for 5-8 days under long-day conditions after stratification for 2 d (means ± SD, *n* = 5-12). (c) Plants were grown under long-day conditions for four weeks. (d) Chlorophyll fluorescence of 4-week-old plants grown on soil under long-day conditions (means ± SD, *n* = 7). The statistical analysis (one way ANOVA with post hoc Holm-Sidak comparisons for WT vs. mutant) indicates significant changes; \**P* ≤ 0.05; \*\*\**P* ≤ 0.001.

### NADPH-dependent thioredoxin reductases backup mitochondrial GR2

Both, NTRA and NTRB, are known to be dual-targeted to the cytosol and mitochondria (Reichheld *et al*., 2005). Based on the observation that the NTR/TRX system efficiently reduces GSSG in the cytosol (Marty *et al*., 2009) we hypothesized that NTRA and NTRB together with mitochondrial TRXo1 and o2 (Yoshida & Hisabori, 2016), or TRXh2 (Meng *et al*., 2010) may provide a functional backup system for mitochondrial GR2. To further support this hypothesis, we also tested mitochondrial TRXs for their ability to reduce GSSG *in vitro* and determined steady-state kinetic parameters for GSSG for TRXh2, TRXo1 and GR2. In conjunction with NTRA recombinant TRXh2 and TRXo1 are both capable of reducing GSSG, albeit with *K_m_* values of only 2 to 4×10^3^ µM, which is nearly 100-fold higher than the *K_m_* of GR2 (13 µM) (Fig. 9). While the efficiency *k_cat_/K_m_* of GR2 approximates the diffusion limit, *k_cat_/K_m_* for both TRXs is about 1000-fold lower. The GSSG reduction activity of both TRXs is independent of which NTR isoform is available for TRX reduction (Fig. S6).

**Fig. 9.**
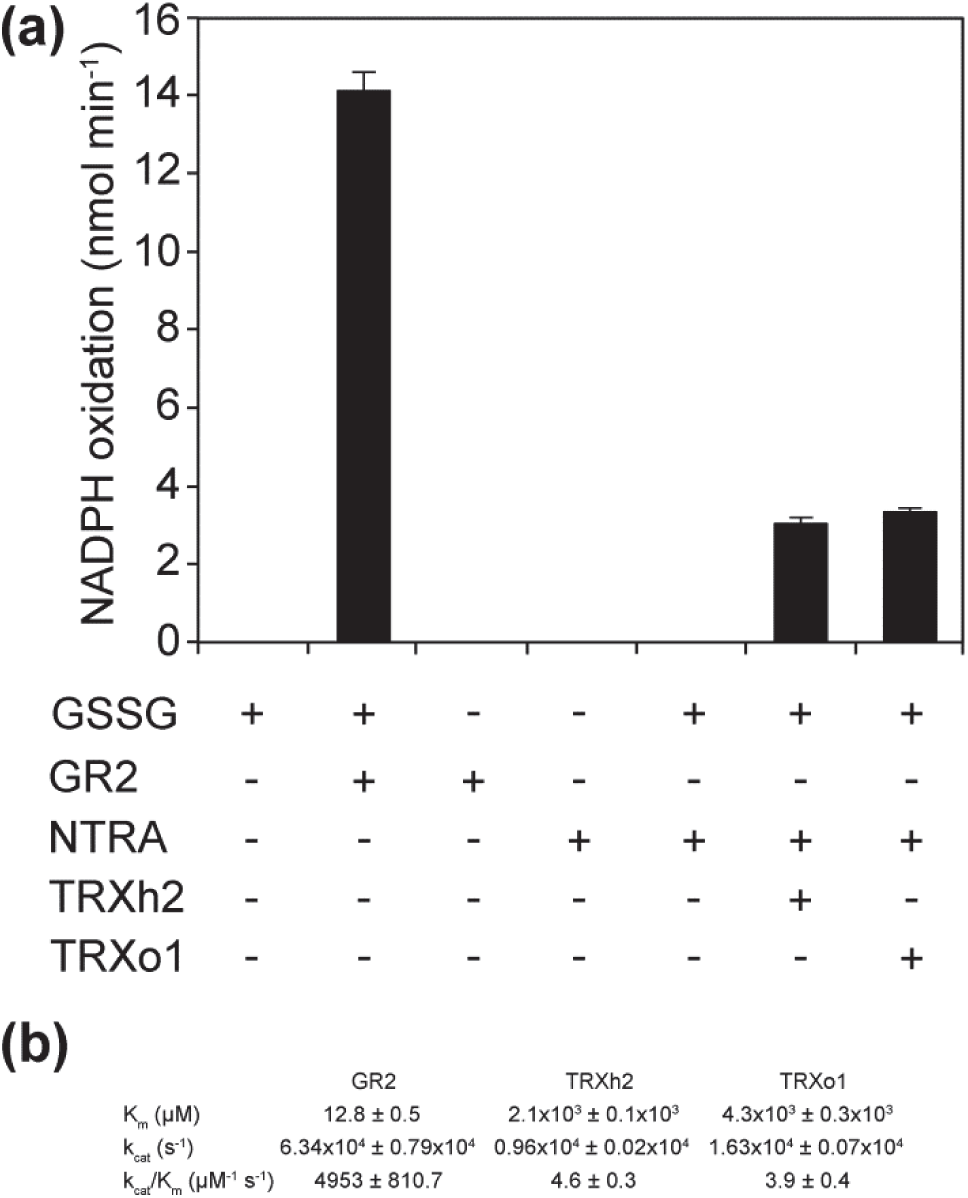
Mitochondrial TRXs in conjunction with NTRA can reduce GSSG *in vitro*. (a) Activity is monitored as NADPH oxidation. Enzymes and substrates were used at the following concentrations: NADPH, 250 µM; GSSG, 1 mM; GR2, 0.01 µM; TRXh2 and TRXo1, 2 µM; NTRA, 1 µM; Means ± SD (*n* = 3). Note that the amount of disulfide reductase protein varied between the assays. (b) Steady-state kinetic parameters of GR2 and TRX-dependent GSSG reduction systems.

To further test the hypothesis that mitochondrial NTRs provide the rescue for lack of mitochondrial GR2, we generated a cross between an *ntra ntrb* double null mutant and a homozygous *gr2* complemented with plastid-targeted GR2 (Fig. S7). From this cross, F_3_ plants expressing GR2 were preselected by Basta^®^ to ensure only GR2 complemented plants to be maintained. Next, plants segregating for *ntrb* only were selected through genotyping, and selfed. 71 tested F_4_ progeny segregated for the mutant allele *ntrb* 23 *NTRB/NTRB : 48 NTRB/ntrb* : 0 *ntrb/ntrb* (χ^2^ = 0.03, *P* > 0.95) indicating that homozygous *ntra ntrb gr2* triple mutants are not viable. The 1:2 segregation of viable plants suggested that the lethal effect of *ntra ntrb gr2* is not gametophytic but rather manifests itself only after fertilization. Consistent with this, viability staining of pollen resulted in nearly 100 % viable pollen (Fig. S8). To further confirm full viability of *ntra ntrb gr2* pollen during pollen tube growth, we backcrossed the *ntra/ntra NTRB/ntrb gr2/gr2 plGR2* plant to WT. This cross resulted in a 1:1 segregation for *ntrb* (χ^2^ = 0.7, *P* = 0.34; Table S3a). Similarly, the reverse cross of WT pollen to an *ntra/ntra NTRB/ntrb gr2/gr2 plGR2* triple mutant resulted in a 1:1 segregation for *ntrb* (χ^2^ = 0.34, *P* = 0.58; Table S3b). These results showed that both pollen and oocytes carrying all three mutant alleles are fully viable.

Developing siliques from self-fertilized *ntra/ntra NTRB/ntrb gr2/gr2 plGR2* plants in contrast contained many aborted seeds (Fig. 10a,b) supporting that the lethal phenotype was established at the diploid stage. However, the overall abortion rate of 13.4 % in siliques collected from ten randomly selected plants was significantly below the expected 25 % (114 aborted seeds out of 852 seeds; χ^2^ = 61.35, *P* << 0.001). A closer inspection of seedlings germinated from these seeds indicated some barely viable seedlings that ceased growth about three weeks after germination and died (Fig. 10c). PCR genotyping confirmed that these seedlings where homozygous for the segregating allele *ntrb* (Fig. 10d,e).

**Fig. 10.**
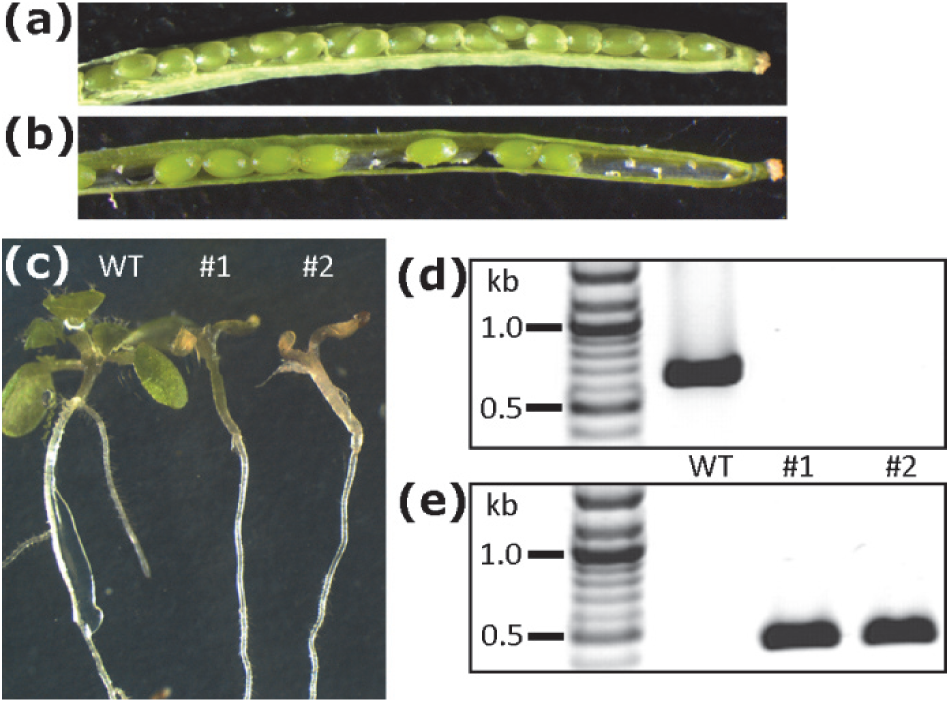
Characterization of *ntra ntrb gr2* triple mutants expressing plastid-targeted GR2 under control of its endogenous promoter. (a) WT silique 20 d after fertilization. (b) Silique from a plant homozygous for both *ntra* and *gr2*, complemented with *plGR2* and heterozygous for *ntrb*. (c) Phenotype of mutant seedlings 18 d after germination. (d, e) Genotyping of seedlings shown in panel c for *NTRB* (d) and the *ntrb* T-DNA insertion (e).

## Discussion

### GR2 is indispensable in plastids

Appropriate detoxification of H_2_O_2_ produced during normal metabolism or in response to environmental stress is mandatory to ensure maintenance of sufficiently reducing conditions for normal metabolism (Moller *et al*., 2007; Waszczak *et al*., 2018). The glutathione-ascorbate cycle has attained much attention and connects H_2_O_2_ detoxification via ascorbate peroxidases ultimately to the local glutathione pool with GSH providing the required electrons (Foyer & Noctor, 2011). The generated GSSG is subsequently reduced back to GSH by GRs with electrons provided by NADPH. While in Arabidopsis GR1 activity in the cytosol and peroxisomes can be substituted to a sufficient extent by the cytosolic NTS to avoid lethality of null mutants (Marty *et al*., 2009), *gr2* null mutants are early embryonic lethal (Bryant *et al*., 2011). Our results unanimously show that GR2 is dual-targeted to plastids and mitochondria but is essential only in plastids.

A shift of *E*_GSH_ towards less negative values can be caused either by depletion of GSH or by increased amounts of GSSG. The GSH-deficient mutant *gsh1* completes embryogenesis up to the seed maturation phase because maternal tissues provide a minimum amount of GSH to the embryo (Cairns *et al*., 2006; Lim *et al*., 2014). In *rml1*, the very low GSH content results in an *E*_GSH_ of about −260 mV (Aller *et al*., 2013). Despite the pronounced shift in *E*_GSH_ the situation is not deleterious *per se*. Partial suppression of the early embryo arrest in *gr2 gsh1* double mutants rather points at the accumulation of GSSG being toxic. The severely restricted ability to build up normal GSH levels initially helps keeping the GSSG concentration low and enables embryos to develop beyond globular stage. However, the arrest at torpedo stage still occurs much earlier than the arrest in *gsh1* during seed maturation. This strongly suggests that even without normal levels of GSH, GSSG quickly accumulates to toxic levels in plastids if no GR2 is present. Plastids apparently have no backup system for GSSG reduction and no (or insufficient) ability to export GSSG to the cytosol for reduction. In developing Arabidopsis embryos, the first chloroplasts start differentiating and greening with the initiation of cotyledons at heart stage (Mansfield & Briarty, 1991). Silencing of GR2 in mature plants indicates that light-dependent reduction of GSSG via plastidic TRXs may not be sufficient to maintain growth. This suggests that, in contrast to the cytosolic NTS, the plastidic FTR/TRX system cannot compensate for the lack of GR activity. This conclusion is consistent with current findings in *Physcomitrella patens* plants lacking organellar GR, where stromal *E*_GSH_ is oxidised but not rescued by active photosynthetic electron transport (Müller-Schüssele *et al*., 2019). Developmental arrest of *gr2* mutants at globular stage before chloroplast differentiation indicates that the accumulation of GSSG is not related to photosynthesis as a possible source of ROS (Foyer & Noctor, 2009), but rather to other basic metabolic processes. Possible processes include the formation of GSSG through reduction of oxidised GRX by GSH (Begas *et al*., 2017) and the reduction of adenosine 5’-phosphosulfate by GSH(Bick *et al*., 1998). In addition, several metabolic pathways in plastids involve oxidation steps in which molecular oxygen is reduced to H_2_O_2_. The reactions include the pyridoxal 5’-phosphate salvage pathway in which the plastidic pyridoxine/pyridoxamine 5’-phosphate oxidase produces one molecule H_2_O_2_ (Sang *et al*., 2011), the oxidation of L-aspartate by L-aspartate oxidase in the early steps of NAD biosynthesis (Katoh *et al*., 2006), and the three-step oxidation of protoporphyinogen IX to protoporphyrin IX by the enzyme protoporphyrinogen oxidase PPOX (Koch *et al*., 2004; Mochizuki *et al*., 2010).

Minute amounts of GR2 activity are necessary to avoid accumulation of GSSG to toxic levels. GR, in general, has been reported to be highly active with a very low K_m_ for GSSG leaving only low nanomolar traces of GSSG (Veech *et al*., 1969). This activity dominates *E*_GSH_ *in vivo* keeping it highly reduced as confirmed by *in vivo* imaging with roGFP-based probes (Marty *et al*., 2009; Schwarzländer *et al*., 2016). Accumulating GSSG may interfere with fundamental plastidic processes, such as transcription, translation and enzymatic functions. For instance, fatty acid biosynthesis is vital in developing oilseed embryos and involves the redox-regulated heteromeric plastidic acetyl-CoA-carboxylase (ACCase) (Ke *et al*., 2000; Sasaki & Nagano, 2004; Bryant *et al*., 2011). The extraordinary importance of plastidic ACCase for embryo development is further supported by the early embryo lethal phenotype of null mutants for BCCP1 (Li *et al*., 2011). One of four ACCase subunits, CTβ (AtCg00500, AccD), is plastid encoded. Thus, interfering with thiol switches regulating the plastidic gene expression machinery (Dietz & Pfannschmidt, 2011) or vital enzyme activities would constitute one possible scenario to account for our observations. Another possible toxic effect of GSSG is the disruption of Fe-S cluster transfer by monothiol GRXs which involves binding of GSH as a cofactor on its backbone (Moseler *et al*., 2015). If present in high concentrations GSSG may interfere with GSH binding and disrupt Fe-S coordination (Berndt *et al*., 2007). Plastids contain two monothiol GRXs of which GRXS16 carries an additional regulatory disulfide that is responsive to GSSG (Zannini *et al*., 2019). GSSG-mediated oxidation of GRXS16 modulates its oxidoreductase function and enables protein glutathionylation. Dissecting the mechanistic cause of embryo lethality is beyond the scope of this work, but promises intriguing novel insights into the role of redox dynamics in organelle biogenesis and early plant development.

### ATM3 and the mitochondrial TRX system safeguard the matrix *E*_GSH_

Direct comparison of cytosol and mitochondrial matrix in plastid complemented *gr2* mutants clearly showed that the readout of roGFP2-Grx1 was shifted to higher ratio values indicating a shift in the local *E*_GSH_ towards less negative values. While roGFP2 in the cytosol of plastid-complemented *gr2* lines is almost fully reduced indicating an *E*_GSH_ of −310 mV or even more negative, roGFP2 in the mitochondrial matrix is clearly partially oxidised. The observed degree of sensor oxidation of ∼50 % would indicate an *E*_GSH_ of −280 mV assuming a matrix pH of 7.0 (−310 mV at pH 7.5). In a solution with estimated 2 mM GSH this point would be reached with approximately 200 nM GSSG. This calculation shows that the amount of GSSG determined by HPLC in plants extracts is a significant overestimation of the amount of GSSG really present in the respective subcellular compartment. This is in line with earlier conclusions on sensor-based subcellular *E*_GSH_ measurements (Meyer & Dick, 2010; Schwarzländer *et al*., 2016).

Although we showed that GSSG accumulation in plastids is causing embryo lethality, plants lacking GR2 in mitochondria have no obvious phenotype under control conditions. Two possible scenarios can be drawn to explain this observation: (*i*) GSSG can be exported to the cytosol to get reduced by GR1, or (*ii*) GSSG is reduced in the matrix by other enzymes. Knock-down mutants with a severely limited ATM3 capacity have been shown earlier to have a less negative *E*_GSH_ in the mitochondrial matrix than WT (Schaedler *et al*., 2014). The pronounced chlorotic phenotype of *gr2 atm3-4* supports the interpretation that ATM3 export GSSG from mitochondria under physiological conditions. This mechanism may be important when NADPH is low, and GR2 activity limited. It should be noted, however, that toxic effects of GSSG in this case may also result from competition with other substrates of ATM3, such as persulfides as required for Fe-S cluster biosynthesis in the cytosol. In the presence of high concentrations of GSSG and with limited transport capacity in *atm3-4* these metabolites may not be exported efficiently.

Dithiol GRXs are capable of catalysing the reduction of GSSG by dihydrolipoamide (Porras *et al*., 2002). In contrast to several non-plant species, Arabidopsis mitochondria contain only the monothiol GRXS15 which was found inactive as an oxidoreductase (Moseler *et al*., 2015). Non-catalysed reduction of GSSG by dihydrolipoamide would be extremely inefficient and unlikely to contribute efficiently to GSSG removal. In contrast, the GSSG reduction activity of NTRs together with mitochondrial TRXs appears high enough to reduce significant amounts of GSSG. Lethality of *gr2 epc-2 ntra ntrb* shows that the situation in the mitochondrial matrix resembles the NTS-based backup of cytosolic GR1 (Marty *et al*., 2009). The presence of multiple backup systems may explain why the lethal effect becomes apparent only after germination. In contrast to *gr2* plastids where direct detection of glutathionylated proteins, protein disulfides and further thiol modifications is not possible due to the minute amount of material in early embryos, proteomic studies in mitochondria lacking GR2 seem feasible. In the future, such measurements may reveal novel metabolic bottlenecks generated from oxidative modification of protein thiols and resulting imbalances in thiol switching.

## Acknowledgements

This work was supported by the Deutsche Forschungsgemeinschaft (DFG) through a project grant (ME1567/3-2), the Emmy-Noether programme (SCHW1719/1-1), the priority program SPP1710 ‘Dynamics of thiol-based redox switches in cellular physiology’ (ME1567/9-1, SCHW1719/7-1, HE 1848/16-1, WI3560/2-1) and a DAAD-Procope exchange grant to AJM and JPR. SAKB was supported through a grant from the HEC Pakistan. Selected aspects of the work in the lab of RH/MW was supported by the DFG within the collaborative research center 1036, TP13. This study is set within the framework of the “Laboratoires d’Excellences (LABEX)” TULIP (ANR-10-LABX-41) and was supported in part by the Centre National de la Recherche Scientifique and by the Agence Nationale de la Recherche ANR-Blanc Cynthiol 12-BSV6-0011.

## Author contributions

AJM, RH and JPR designed research; LM, DB, CM, SAKB, AM, MS, BZ, CR, and MW performed research; JB contributed plant lines; LM, DB, CM, MS, SJMS, JPR and AJM analyzed data and SJMS, MS and AJM wrote the manuscript with assistance from all co-authors.

## Supporting Information

**Fig. S1** Schematic representation of constructs used to confirm signal peptide functionality and for compartment-specific complementation of *gr2* mutants.

**Fig. S2** Characterization of transgenic Arabidopsis lines over-expressing plastid-targeted GR2.

**Fig. S3** Protein gel blot analysis of GR1 and GR2 in mitochondrial preparations of WT and plastid-complemented *gr2* (*epc-2*).

**Fig. S4** Characterization of *gr2 rml1* double mutants.

**Fig. S5** High expression of SHMT_TP_-roGFP2-Grx1 results in incomplete mitochondrial targeting.

**Fig. S6** Mitochondrial thioredoxins reduce GSSG *in vitro* with electrons provided by NTRA and NTRB with similar efficiencies.

**Fig. S7** Crossing scheme for generation of *gr2 ntra ntrb*.

**Fig. S8** Viability stain of pollen from *ntra*^−/−^ *gr2*^−/−^ *NTRB/ntrb plGR2* plants.

**Table S1** Oligonucleotides used in this study.

**Table S2** Genetic complementation of the *gr2* mutant with compartment-specific *GR2* constructs.

**Table S3** Reciprocal cross between *ntra/ntra* NTRB/*ntrb gr2/gr2 plGR2* and WT.

**Methods S1** Antibody production and gel blot analysis.

**Methods S2** Immunogold labelling and electron microscopy

