## Supporting Information for "Arabidopsis glutathione reductase 2 is indispensable in plastids, while mitochondrial glutathione is safeguarded by additional reduction and transport systems"

**The following Supporting Information is available for this article:**

**Fig. S1** Schematic representation of constructs used to confirm signal peptide functionality and for compartment-specific complementation of *gr2* mutants.

**Fig. S2** Characterization of transgenic Arabidopsis lines over-expressing plastid-targeted GR2.

**Fig. S3** Protein gel blot analysis of GR1 and GR2 in mitochondrial preparations of WT and plastid-complemented *gr2* (*epc-2*).

**Fig. S4** Characterization of *gr2 rml1* double mutants.

**Fig. S5** High expression of $SHMT_{TP}$-roGFP2-Grx1 results in incomplete mitochondrial targeting.

**Fig. S6** Mitochondrial thioredoxins reduce GSSG *in vitro* with electrons provided by NTRA and NTRB with similar efficiencies.

**Fig. S7** Crossing scheme for generation of *gr2 ntra ntrb.*

**Fig. S8** Viability stain of pollen from *ntra*$^{-/-}$ *gr2*$^{-/-}$ *NTRB/ntrb plGR2* plants.

**Table S1** Oligonucleotides used in this study.

**Table S2** Genetic complementation of the *gr2* mutant with compartment-specific *GR2* constructs.

**Table S3** Reciprocal cross between *ntra*/*ntra* NTRB/*ntrb gr2/gr2 plGR2* and WT.

**Methods S1** Antibody production and gel blot analysis.

**Methods S2** Immunogold labelling and electron microscopy

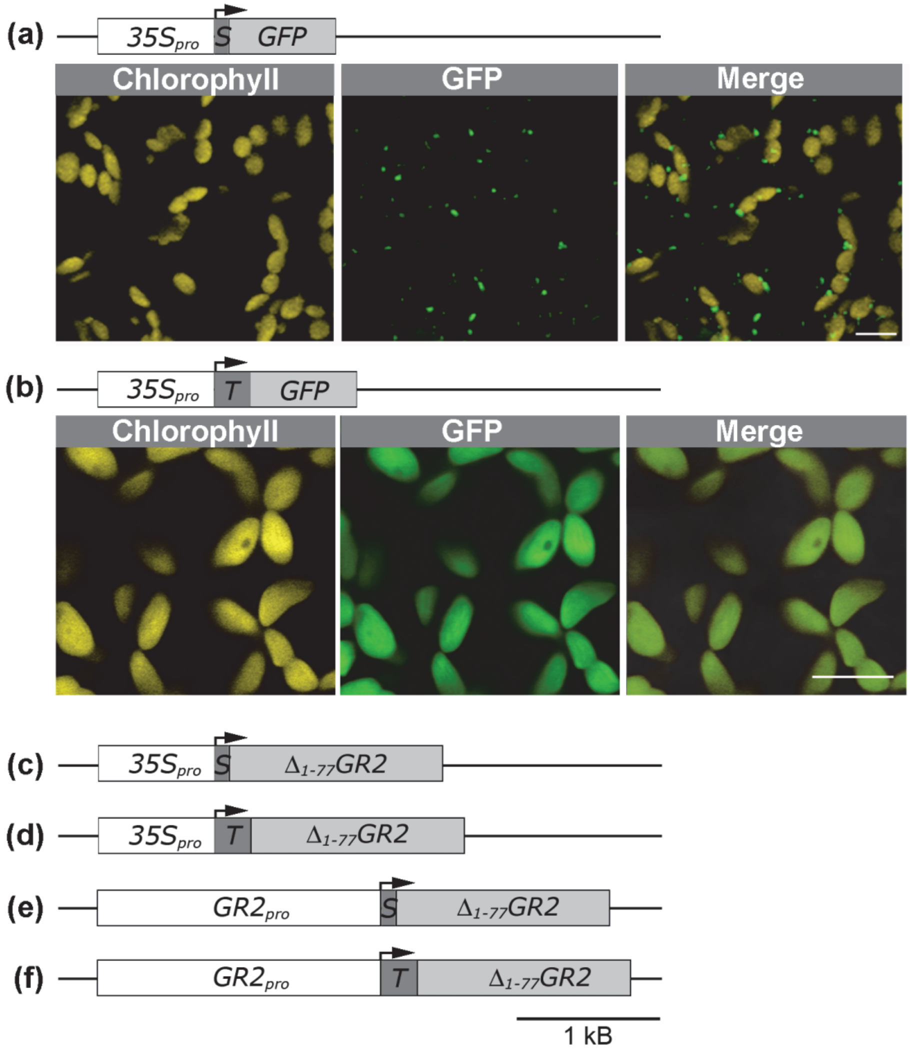


**Fig. S1** Schematic representation of constructs used to confirm signal peptide functionality (a, b) and for compartment-specific complementation of *gr2* mutants (c-f). White boxes indicate promoter elements, dark grey boxes indicate the signal peptides used (S: $SHMT_{TP}$; T: $TK_{TP}$), light grey boxes represent GFP reporter gene or truncated versions of *GR2* ($\Delta_{1-77}$*GR2*) lacking the endogenous target peptide, respectively. GFP constructs (a, b) were cloned in pBinAR, $\Delta_{1-77}$*GR2* constructs used for complementation of *gr2* were cloned in pBarA. Promoter elements controlling the transgene were either *CaMV 35S* promoter ($35S_{pro}$) (a-d), or the endogenous *GR2* promoter from Arabidopsis ($GR2_{pro}$) (e, f). Bars in panels a and b, 10 µm.

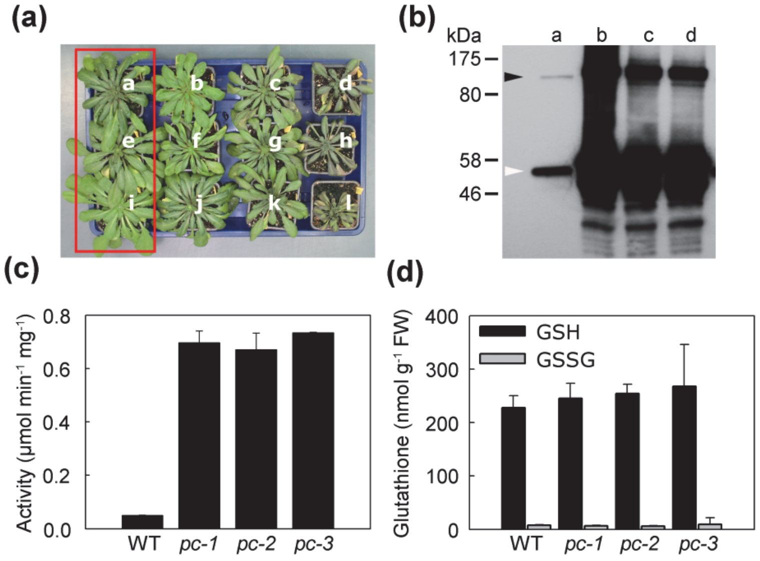


**Fig. S2** Characterization of transgenic Arabidopsis lines over-expressing plastid-targeted GR2. (a) Phenotypes of Basta®-selected T1 *gr2* mutants transformed with a *35S$_{pro}$:TK$_{TP}$-$\Delta_{1\text{-}77}$GR2* construct. Genotyping of complemented plants for the *gr2* locus gave the following results ('+' stands for the WT allele and '-' for the defective T-DNA insertion allele): a: -/- (*pc-1*), b: +/-, c: -/-, d: -/-, e: -/- (*pc-2*), f: -/-, g: +/-, h: +/-, i: -/- (*pc-3*), j: +/+, k: +/-, l: -/-. Three homozygous *gr2* plants carrying the complement and lacking an obvious phenotype were selected for further analysis (lines *pc-1* to *pc-3*, red box). (b) Protein gel blot analysis of GR2 in WT (lane a), *pc-1* (lane b), *pc-2* (lane c) and *pc-3* (lane d). Proteins detected with GR2 antiserum in WT had the size of ~110 kDa (black arrow) and ~53 kDa (white arrow). (c) GR activity of total protein extracts of WT, and *pc-1*, *pc-2*, *pc-3* lines. Values show means ± SD of three T2 plants. (d) Content of reduced glutathione (GSH) and glutathione disulfide (GSSG) in leaf tissue of WT and transgenic lines. Means ± SD of six independent T2 plants for each line are shown.

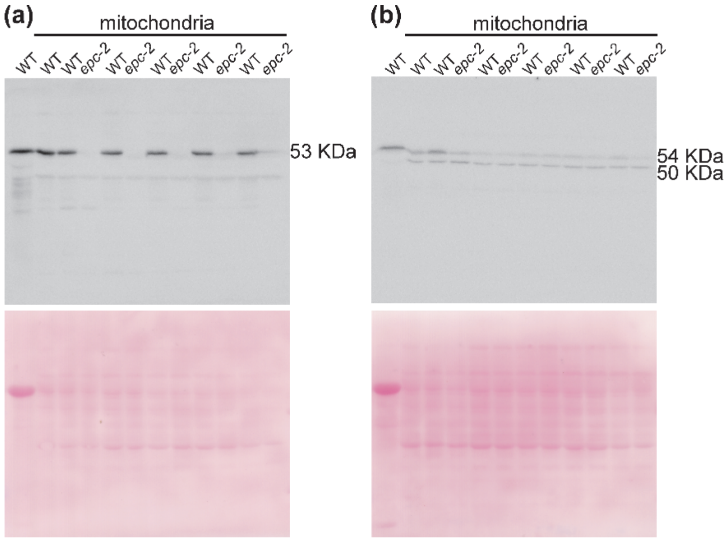


**Fig. S3** Protein gel blot analysis of GR1 and GR2 in mitochondrial preparations of WT and plastid-complemented *gr2* (*epc-2*). (a) Hybridization of membrane with anti-GR2. Mitochondrial preparations from *epc-2* contain no GR2. (b) Hybridization of membrane with anti-GR1 resulted in labelling of all mitochondrial preparations from WT and *epc-2*. Labelling always occurred in two bands with a molecular weight of about 54 and 50 kDa. Lack of the second band with an apparent molecular weight of 50 kDa in whole leaf extracts of WT plants (lane 1) indicates that this band results from enrichment of mitochondrial proteins in the organelle preparations and cross reaction of the antibody with a protein other than GR1. Because of close evolutionary relation to GR1 it is likely that the band represents two mitochondrial lipoamide dehydrogenases (MTLPD1 and MTLPD2), which indeed both have a predicted molecular weight of 49.9 kDa.

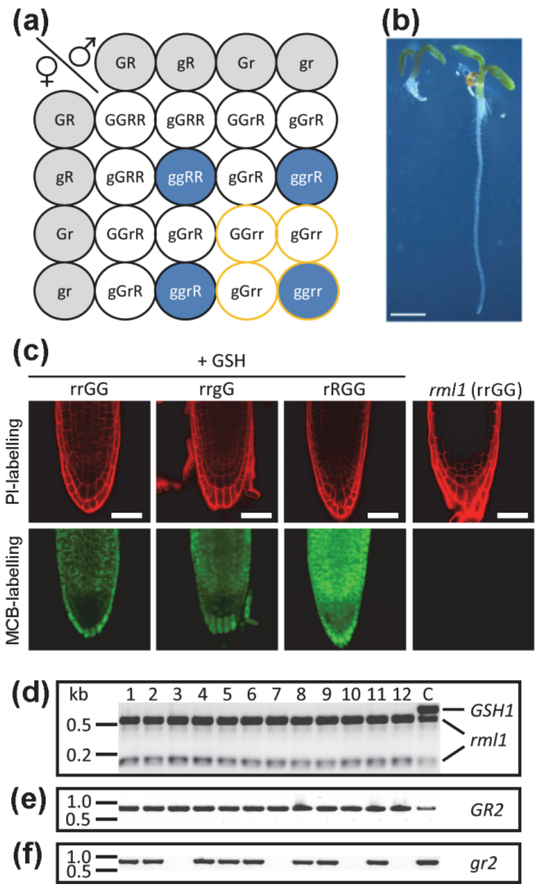


**Fig. S4** Characterization of *gr2 rml1* double mutants. (a) Crossing scheme with expected genotypes for *GSH1* and *GR2* loci. Gametes are depicted in grey. Progeny homozygous for *gr2* are shown in blue and homozygous *rml1* mutants are shown with a yellow circle. R: *GSH1*, r: *rml1*, G: *GR2*, g: *gr2*. (b) The progeny from a *gr2*$^{+/-}$ *rml1*$^{+/-}$ double heterozygous plant segregates with phenotypes indicative of *rml1* (left) and WT (right). Bar, 1 cm. (c) *In situ* labelling of GSH in root tips of *rml1*-like progeny (rrGG and rrgG) and a seedling displaying a WT phenotype (rRGG) 12 days after transfer to plates supplemented with 1 mM GSH. A root tip of *rml1* germinated on GSH-free plates is shown for comparison. Roots were labelled with 100 µM MCB (green) and 50 µM PI (red). Bars, 50 µm. (d-f) Genotyping of seedlings with a typical *rml1*-like phenotype for the *rml1* mutation (d), the *GR2* WT allele (e), and the *gr2* T-DNA insertion (f).

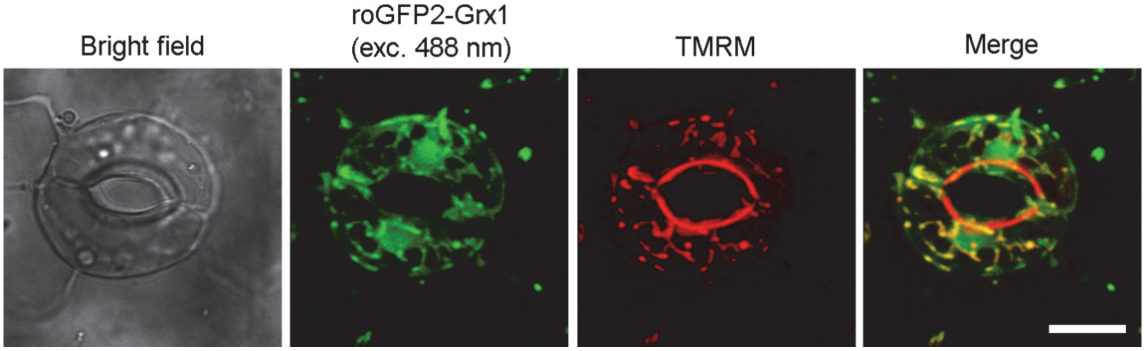


**Fig. S5** High expression of SHMT$_{TP}$-roGFP2-Grx1 results in incomplete mitochondrial targeting. Co-localization with the mitochondrial marker tetramethylrhodamine methyl ester (TMRM) indicates the roGFP-labelled mitochondria in the merge image in yellow. The outer cuticular ledges are strongly labelled appear red because the lipophilic TMRM incorporates in the cuticular waxes. Bar, 10 µm.

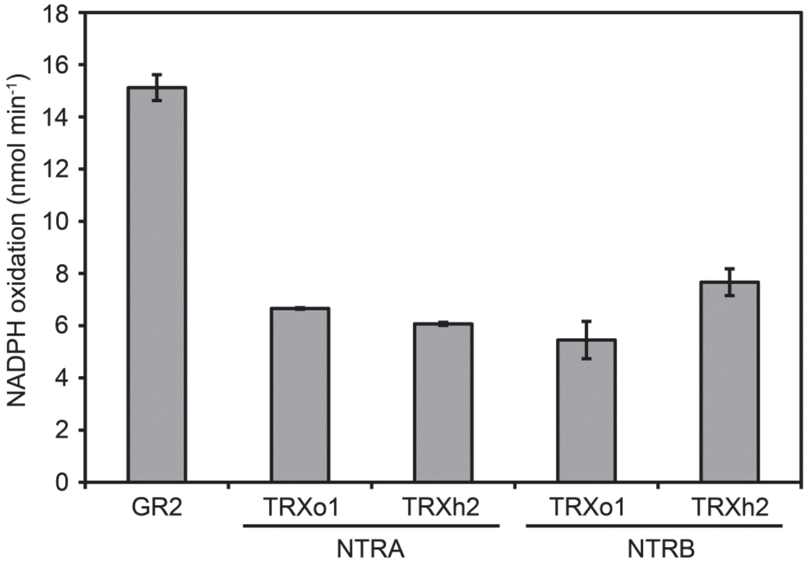


**Fig. S6** Mitochondrial thioredoxins reduce GSSG *in vitro* with electrons provided by NTRA and NTRB with similar efficiencies. GSSG reduction activity is monitored as NADPH oxidation. Enzymes and substrates were used at the following concentrations: NADPH: 250 µM; GSSG: 1 mM; GR2: 0.01 µM; TRXh2 and TRXo1: 2 µM; NTRA and NTRB: 1 µM; Means ± SD (*n* = 3).

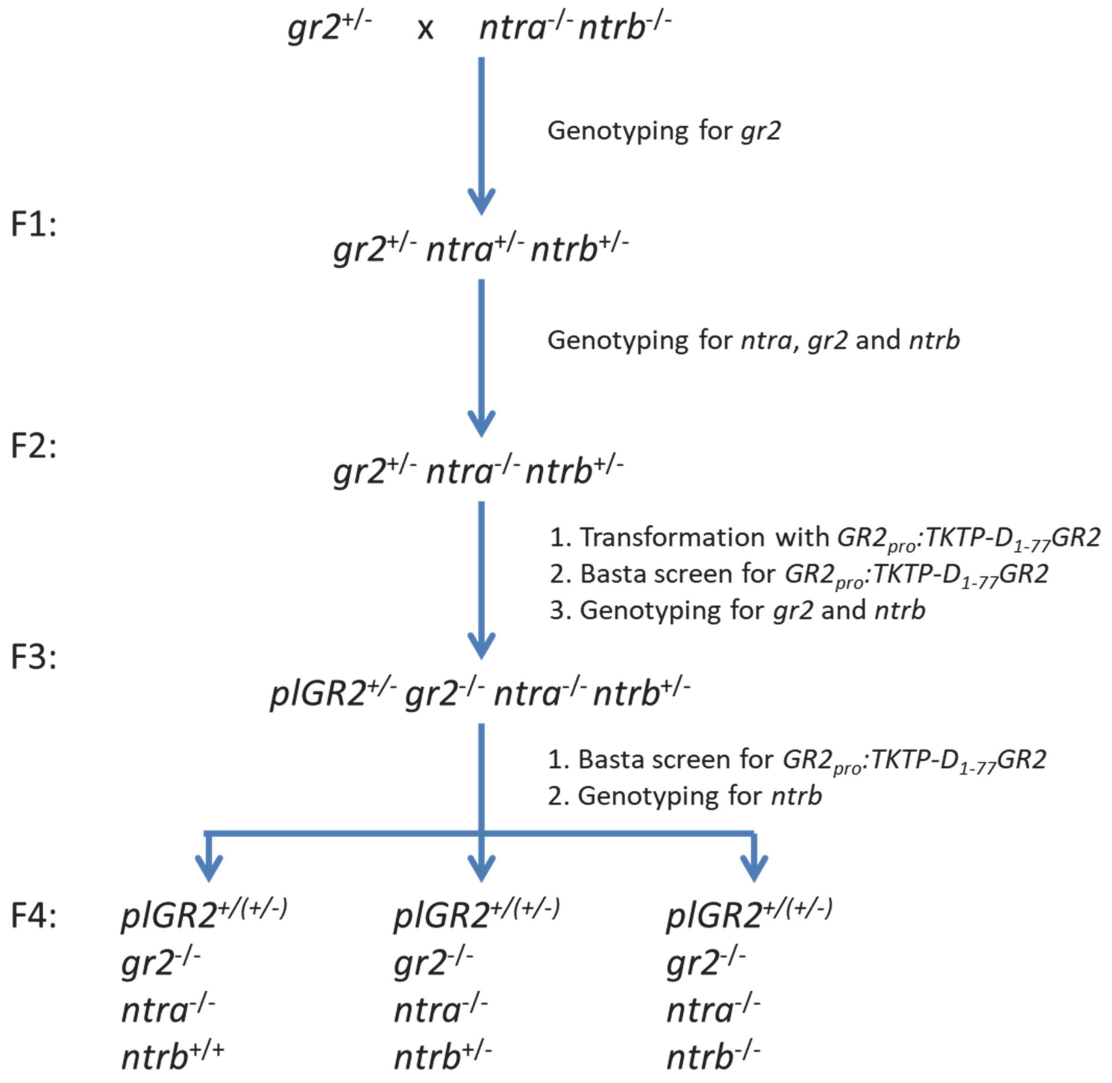


**Fig. S7** Crossing scheme for generation of *gr2 ntra ntrb.* For the F1, F2 and F3 only genotypes selected for further breeding are shown. *plGR2* refers to the plastid-targeted *GR2* construct obtained through transformation with $GR2_{pro}$:*TKTP*-$D_{1\text{-}77}$*GR2*.

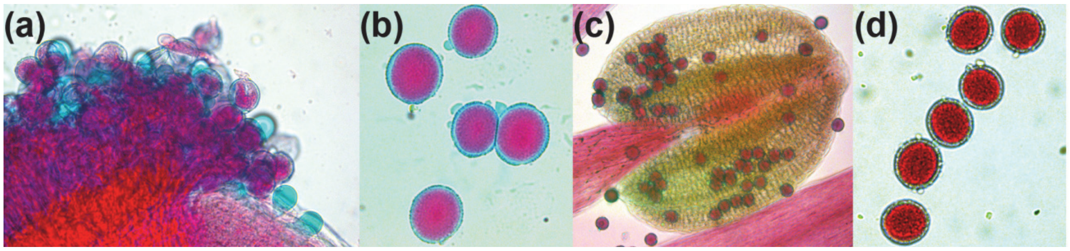


**Fig. S8** Viability stain of pollen from *ntra*$^{-/-}$ *gr2*$^{-/-}$ *NTRB/ntrb plGR2* plants. (a, b) Staining of wild-type pollen results in red colouring of the cytosol while the pollen coat is stained in blue (b). Only germinated pollen on the pistil show exclusive blue colouring (a) which would also be expected for dead pollen if present. (c, d) Pollen from *ntra*$^{-/-}$ *gr2*$^{-/-}$ *NTRb/ntrb plGR2* are evenly stained blue-red without any indication of segregating dead pollen.

**Table S1** Oligonucleotides used in this study.

| | |
|---|---|
| GR2 forward | 5'-GCTACCCTTTCAGGACTTCCAGACC-3' |
| GR2 reverse | 5'-CACAATGTTCTCCTGCAAACATGC-3' |
| T-DNA left border primer | 5'-GACCGCTTGCTGCAACTCTCTCAGG-3' |
| NTRA forward | 5'-GCCGTCGACATGGAAACTC-3' |
| NTRA reverse | 5'-GCTCTCTGCTGCATAATCTTAG-3' |
| NTRB forward | 5'-GAGCGTCTAAGATTATGCAGC-3' ? |
| NTRB reverse | 5'-GATCTCTCTACTAAGCATGGA-3' |
| RML1 forward | 5'-GCCAATGCTCTCACCCTAAA-3' |
| RML1 reverse | 5'-GGCAATGGTTAGTCAAAATCG-3' |
| ATM3 forward | 5'-ATTGCTACACTTGCGGGAGATGC-3' |
| ATM3 reverse | 5'-GATGGTGAGTTATCTGAGAGG-3' |
| GSH1 forward | 5'-TGTTTCGGGTGGCGTGAG-3' |
| GSH1 reverse | 5'-GCTTTCCCTGGTCAACAA-3' |
| $SHMT_{TP}$ forward | 5'-GGTACCATGGCCATGGCTCTTCG-3' |
| $SHMT_{TP}$ reverse | 5'-GGATCCGGGACAGCTTCACTGGGC-3' |
| roGFP2 for SHMT/$TK_{TP}$ forward | 5'-GGATCCCATGGTGAGCAAGGGCGAG-3' |
| roGFP2 for $SHMT_{TP}$/$TK_{TP}$ reverse | 5'-GTCGACTTACTTGTACAGCTCGTCC-3' |
| GR2 for $SHMT_{TP}$/$TK_{TP}$ forward | 5'-GGATCCTAATGGAGCTGAATCAGACCG-3' |
| GR2 for $SHMT_{TP}$/$TK_{TP}$ reverse | 5'-GTCGACCTACACCCCAGCAGCTG-3' |
| GR2 promoter forward | 5'-GAATTCGATGTTTGTGTAGTATGTTTGTTC-3' |
| GR2 promoter reverse | 5'- GGTACCTTCGATAATTCTACTTGTGGCAT-3' |
| Human Grx1 forward | 5'-AAGAATTCGCTCAAGAGTTTGTGAACTGCAAAATCCAG-3' |
| Human Grx1 reverse | 5'-CCATCGATTTACTGCAGAGCTCCAATCTGCTTTAG-3' |
| roGFP2-Grx1 forward | 5'-AAGGATCCGGTGAGCAAGGGCGAGGAGC-3' |
| roGFP2-Grx1 reverse | 5'- TATGTCGACTTACTGCAGAGCTCCAATCTGCTTTA-3' |

**Table S2** Genetic complementation of the *gr2* mutant with compartment-specific *GR2* constructs.

| Complementation construct | Genotyped plants[a] | *GR2/GR2* | *GR2/gr2* | *gr2/gr2* |
|---|---|---|---|---|
| $35S_{pro}$:$SHMT_{TP}$-$\Delta_{1\text{-}77}$GR2 | 60 | 43 % (26) | 57 % (34) | 0 % |
| $35S_{pro}$:$TK_{TP}$-$\Delta_{1\text{-}77}$GR2 | 60 | 23 % (14) | 55 % (33) | 22 % (13) |
| $GR2_{pro}$:$SHMT_{TP}$-$\Delta_{1\text{-}77}$GR2 | 48 | 56 % (27) | 44 % (21) | 0 % |
| $GR2_{pro}$:$TK_{TP}$-$\Delta_{1\text{-}77}$GR2 | 69 | 28 % (19) | 62 % (43) | 10 % (7) |

[a] T1 plants were genotyped five weeks after germination (number of plants with the respective genotype is given in parenthesis).

**Table S3** Reciprocal cross between *ntra/ntra* NTRB/*ntrb gr2/gr2 plGR2* and WT.

| **(a)** | Female parent | X | Male parent | Progeny genotype | |
|---|---|---|---|---|---|
| | WT | X | *gr2/gr2* *plGR2/plGR2* *ntra/ntra* *NTRB/ntrb* | *GR2/gr2* *plGR2/-* *NTRA/ntra* *NTRB/ntrb* | *GR2/gr2* *plGR2/-* *NTRA/ntra* *NTRB/NTRB* |
| | Observed frequency | | | 39 | 32 |
| | $\chi^2$ for 1:1 segregation of *ntrb* | | | 0.7 ($P$ = 0.34) | |

| **(b)** | Female parent | X | Male parent | Progeny genotype | |
|---|---|---|---|---|---|
| | WT | X | *gr2/gr2* *plGR2/plGR2* *ntra/ntra* *NTRB/ntrb* | *GR2/gr2* *plGR2/-* *NTRA/ntra* *NTRB/ntrb* | *GR2/gr2* *plGR2/-* *NTRA/ntra* *NTRB/NTRB* |
| | Observed frequency | | | 5 | 7 |
| | $\chi^2$ for 1:1 segregation of *ntrb* | | | 0.34 ($P$ = 0.58) | |

### Methods S1

#### Antibody production and gel blot analysis

Recombinant GR2 proteins lacking their endogenous target peptide were purified by $Ni^{2+}$ affinity chromatography and used as antigens. The purity of the protein was confirmed by SDS-PAGE gel stained with Coomassie. After size exclusion chromatography fractions containing the GR2 protein were concentrated using a Vivaspin 2-Maximunspin column. 200 µg GR2 protein in a volume of 300 µl was mixed with 300 µl complete Freund's adjuvant and used for rabbit injection. Before the first immunization, blood was taken as pre-immune serum. Six weeks after the first immunization the first boost-step with 300 µl incomplete Freund's adjuvant mixed with 300 µl of 200 µg pure protein was performed. Two weeks after the first boost, antibody production was monitored. After confirmation that antibody production was successful after the first boost, the final bleed was taken. After 5 h at room temperature the coagulated blood was removed by centrifugation and sodium azide was added to the antiserum to a final concentration 0.02 %. Aliquots of the antiserum were stored at -80 °C and 4 °C.

For western blot analysis 15 µg of extracted leaf proteins or 20 µg of mitochondrial proteins were separated by discontinuous SDS-PAGE in Mini-Protean II system (Bio-Rad). Proteins were transferred to nitrocellulose or PVDF membrane using the Mini Trans-Blot system (Bio-Rad) or the Criterion Blotter (Bio-Rad). Subsequently the membrane was blocked for 2–12 h at room temperature with 5 % milk powder in TBS (20 mM Tris-HCl, 137 mM NaCl, pH 7.6) supplemented with 0.1 % v/v Tween 20. The membrane was washed three times with 0.1 % Tween in TBS and incubated with 1:5,000 (for α-PRXII F), 1:2,500 (for α-GR1) or 1:2,000 (for α-GR2) diluted primary antiserum and 0.1 % Tween for 2 h. The membrane was washed three times for 5 min with 0.1 % Tween in TBS. For detection of GR2 the membrane was incubated with either Antirabbit IgG-Alkaline Phosphatase Conjugate or ImmunoPure goat anti-rabbit IgG horseradish peroxidase conjugated antibody (Thermo Scientific) as secondary antibodies. Antirabbit IgG-Alkaline Phosphatase Conjugate (1:10,000 in 0.5 % milk powder, 0.1 % Tween in TBS) was applied for 1 h and washed twice with AP buffer (100 mM Tris-HCl pH 9.5, 100 mM NaCl, 5 mM $MgCl_2$) for 5 min. Alkaline phosphatase reaction was done by adding the substrates nitroblue tetrazolium and 5-bromo-5-chloro-3-indolyl phosphate.

ImmunoPure goat anti-rabbit IgG horseradish peroxidase conjugated antibody (1:20,000 in 0.5 % milk powder, 0.1 % Tween in TBS) was applied and incubated on the membrane for 1 hour. After washing the membrane six times for 5 min with 0.1 % Tween in TBS protein–antibody complexes were visualized by using the Pierce ECL Western Blotting or SuperSignal West Femto Substrate (Thermo Scientific). Chemiluminescence was detected with the MF-ChemiBIS 2.0 imaging system (Biostep).

### Methods S2

#### Immunogold labelling and electron microscopy

Small leaf samples (1.5 $mm^2$) were cut in a drop of 2.5 % paraformaldehyde/0.5 % glutardialdehyde in 0.06 M Sørensen phosphate buffer (pH 7.2) and fixed for 90 min at room temperature (RT). After fixation samples were rinsed in buffer (four times 15 min each) and then dehydrated in increasing concentrations of acetone (50 %, 70 %, and 90 %) twice for 10 min each. Subsequently, specimens were gradually infiltrated with increasing concentrations of LR-White resin (30 %, 60 % and 100 %; London Resin Company Ltd.) mixed with acetone (90 %) for a minimum of 3 h per step. Samples were finally embedded in pure, fresh LR-White resin and polymerized at 50 °C for 48 h in small plastic containers under anaerobic conditions. Immunogold labelling of GR2 was done with ultrathin sections (80 nm) on nickel grids with the automated immunogold labelling system Leica EM IGL (Leica Microsystems). The ideal dilution of the primary and secondary antibodies was determined in preliminary studies by evaluating the labelling density after a series of labelling experiments. The final dilution of the primary and secondary antibody used in this study showed a minimum of background labelling outside the sample with a maximum of specific labelling inside the sample. For cytohistochemical analysis samples were blocked with 5 % bovine serum albumine (BSA) in tris-buffered saline (TBS, pH 7.4) for 20 min at RT. Samples were then treated with the primary antibody against GR diluted 1:100 in TBS containing 1 % BSA for 2 h at RT. After a short rinse in TBS (three times 5 min) the samples were incubated with a 10 nm gold-conjugated secondary antibody (goat anti rabbit IgG, British BioCell International) diluted 1:100 in TBS for 90 min at RT. After a short wash in TBS (three times 5 min) and distilled water (two times 5 min) labelled grids were post-stained with 1 % uranyl-acetate dissolved in Aqua bidest. (15 s) and observed in a Philips CM10 transmission electron microscope (TEM).

Several negative controls were made to confirm the specificity of the immunogold procedure. Negative controls were treated either with: (1) gold conjugated secondary antibody (goat anti rabbit IgG) without prior incubation of the section with the primary antibody, or (2) non-specific secondary antibody (goat anti mouse IgG). Gold particles were virtually absent on these sections (data not shown).

Micrographs of randomly photographed immunogold labelled sections of mesophyll cells were digitized and gold particles were counted automatically using the software package Cell D with the particle analysis tool (Olympus). For statistical evaluation a minimum of 60 sectioned cell structures of at least 15 different cells throughout the block were analysed for gold particle density. The obtained data were statistically evaluated using Statistica (Stat-Soft) and presented as the number of gold particles per $\mu m^2$.
